# Ser/Thr kinase-dependent phosphorylation of the peptidoglycan hydrolase CwlA controls its export and modulates cell division in *Clostridioides difficile*

**DOI:** 10.1101/2020.10.29.360313

**Authors:** Transito Garcia-Garcia, Sandrine Poncet, Elodie Cuenot, Thibaut Douché, Quentin Giai Gianetto, Johann Peltier, Pascal Courtin, Marie-Pierre Chapot-Chartier, Mariette Matondo, Bruno Dupuy, Thomas Candela, Isabelle Martin-Verstraete

## Abstract

Cell growth and division require a balance between synthesis and hydrolysis of the peptidoglycan (PG). Inhibition of PG synthesis or uncontrolled PG hydrolysis can be lethal for the cells, making it imperative to control peptidoglycan hydrolase (PGH) activity. The serine/threonine kinases (STKs) of the Hanks family control cell division and envelope homeostasis, but only a few kinase substrates and associated molecular mechanisms have been identified. In this work, we identified CwlA as the first STK-PrkC substrate in the human pathogen *Clostridiodes difficile* and showed that CwlA is an endopeptidase involved in daughter cell separation. We demonstrated that PrkC-dependent phosphorylation inhibits CwlA export, therefore controlling the hydrolytic activity in the cell wall. High level of CwlA at the cell surface led to cell elongation, whereas low level caused cell separation defects. We thus provided evidence that the STK signaling pathway regulates PGH homeostasis to precisely control PG hydrolysis during cell division.

## Introduction

PG, the major component of the bacterial cell wall (CW) protects bacteria from environmental stresses and contributes to virulence and antibiotic resistance (Scheffers & Pinho, 2005). PG is classically made of glycan chains of alternating N-acetylglucosamine (NAG) and N-acetylmuramic acid (NAM) cross-linked by short stem peptides attached to NAM (Vollmer, Blanot, et al., 2008). PG remodeling requires three types of hydrolases (PGH, or autolysins): (i) glycosidases hydrolyze the glycosidic linkages, (ii) amidases cleave the amide bond between NAM and L-alanine residue and (iii) endopeptidases cleave amide bonds between amino acids within the PG peptidic chains (Vermassen et al., 2019) (**Fig 1a**). PGHs are involved in fundamental aspects of bacterial physiology such as CW expansion, turnover and recycling during growth, cell separation and autolysis (Smith et al., 2000; van Heijenoort, 2011; Vollmer, Joris, et al., 2008). Different strategies are employed by bacteria to coordinate PG synthesis and degradation. Typical regulation occurs through a signal, such as iron concentration that regulates the transcription of autolysin-encoding gene *isdP* in *Staphylococcus lugdunensis* (Farrand et al., 2015) or through two component systems such as WalK-WalR that monitors the expression of the *lytE* and *cwlO* genes encoding endopeptidases in *Bacillus subtilis* by sensing and responding to their cleavage products (Dobihal et al., 2019). PGH activity can also be directly regulated, for instance through the control of their cell surface localization. The FtsEX complex associated with the divisome recruits and activates *Streptococcus pneumoniae* PcsB (Bartual et al., 2014; Rued et al., 2019; Sham et al., 2011) and *B. subtilis* CwlO (Meisner et al., 2013) hydrolases involved in cell division and elongation. In *Escherichia coli*, the interaction of FtsEX with EnvC activates cell-separation amidases (Meisner et al., 2013; Uehara et al., 2010). Alternatively, proteolysis modulates *E. coli* Meps (Singh et al., 2015) and *Mycobacterium tuberculosis* RipA (Botella et al., 2017) hydrolase activities. Post-translational modifications also provide an efficient and fine tunable way to regulate PGH activity. The tyrosine kinase CspD phosphorylates LytA, a pneumococcal autolysin enhancing its amidase activity (Standish et al., 2014), whereas *O*-glycosylation of the *Lactobacillus plantarun N*-acetylglucosaminidase decreases its activity (Rolain et al., 2013). The synthesis or activity of several key enzymes of PG metabolism is also regulated by Hanks-type serine/threonine kinase (STKs). In all firmicutes, a trans-membrane kinase with an extracellular domain containing penicillin-binding and STK associated (PASTA) repeats is present. PASTA domains interact with β-lactam antibiotics (Maestro et al., 2011), non-crosslinked PG fragments (muropeptides) (Squeglia et al., 2011) and lipid II (Hardt et al., 2017; Kaur et al., 2019). Inactivation of genes encoding PASTA-STKs is associated with cell division defects, changes in CW metabolism and modified susceptibility to CW-targeting antibiotics (Manuse et al., 2016; Pensinger et al., 2018; Pereira et al., 2011). In *S. pneumoniae*, STK allows localization of the LytB PGH at the septum through their PASTA repeats (Zucchini et al., 2018). However, only a few STK substrates have been identified so far.

**Figure 1.**
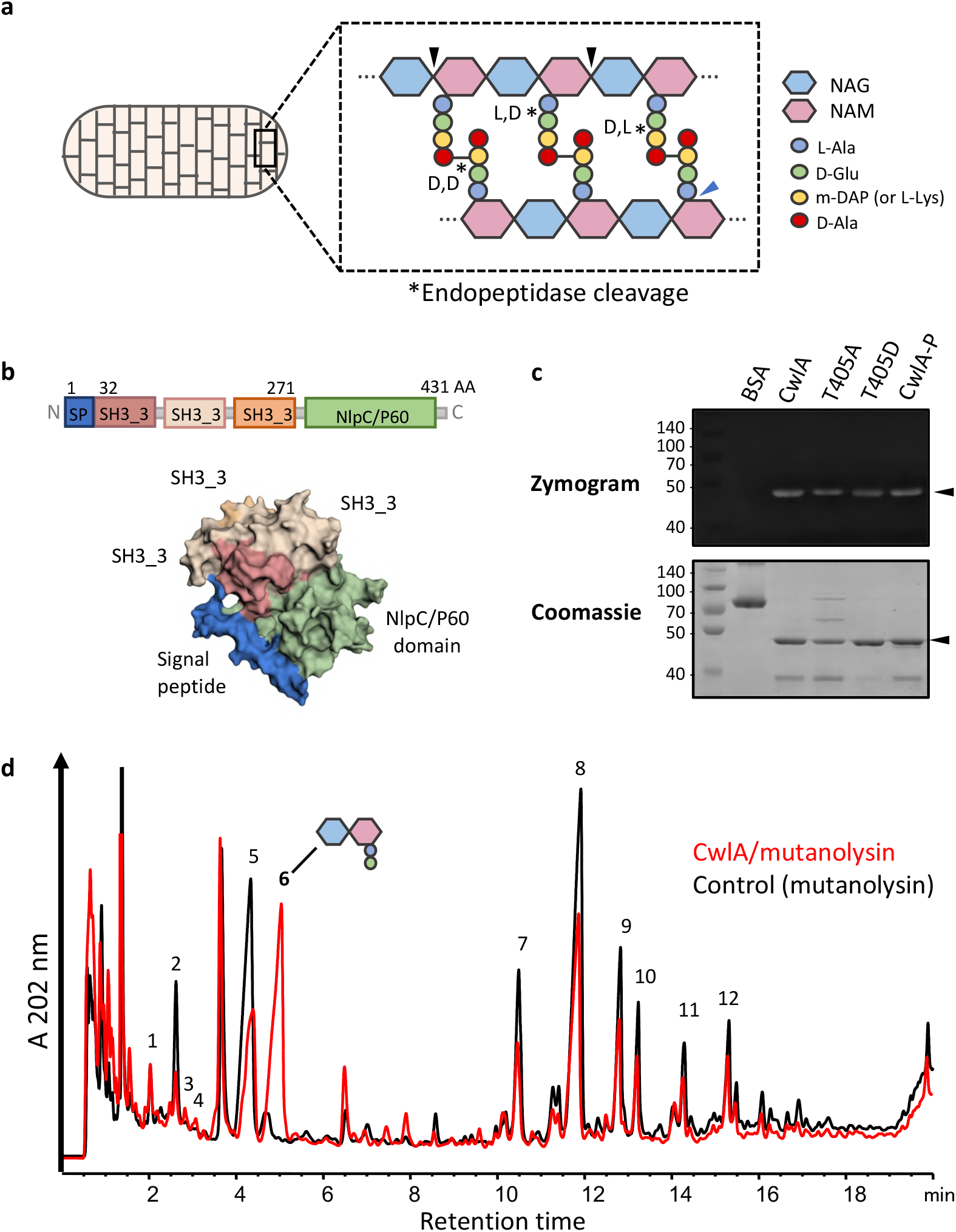
CwlA is γ-D-Glu-mDAP-endopeptidase. **a,** Schematic representation of peptidoglycan sacculus, a polymer of β-(1,4)-linked *N*-acetylglucosamine (NAG, blue) and *N*-acetylmuramic acid (NAM, pink) glycan strands cross-linked by short peptide chains. Enzymes that hydrolyze peptidoglycan are classified as glycosidases (black arrow), amidases (blue arrow) and endopeptidases (*) depending on where they cleave. **b**, Structural prediction of CwlA colored by domains. Signal peptide in blue (1-31 amino acids), three SH3_3 domains (also named SH3b, pfam08239) in pink (32-95), light pink (114-178) and orange (207-271), and the catalytic domain NlpC/P60 (pfam00877) in green. Structural prediction was generated with Phyre 2.0 based on the alignment with <u>c6biqA</u>, <u>c3npfB</u> and <u>c3h41A</u>. Highlighted domains were performed with EzMol 2.1 **c**, Detection by zymogram of hydrolytic activities of purified proteins: BSA (negative control), CwlA, CwlA-T405A (T405A), CwlA-T405D (T405D) and PrkC-dependent phosphorylation of CwlA (CwlA-P). The gel contained 1 mg/ml of purified PG as a substrate. Arrows indicate the positions of the hydrolytic bands. **d**, RP-HPLC separation of muropeptides released from *C. difficile* PG after incubation with mutanolysin (control, black) or His_6_-CwlA followed by mutanolysin (red). The proposed structures deduced from MS analysis are indicated in Table 1. Peak 6 contains deacetylated disaccharide dipeptide (NAG(deAc)-NAM-L-Ala-D-Glu) resulting from CwlA cleavage of PG stem peptides before mutanolysin digestion.

*Clostridioides difficile* contains two STKs of the Hanks family, the PASTA-kinase, PrkC, and a second kinase, CD2148, and one gene encoding a PP2C-type phosphatase (STP), which dephosphorylates the Hanks-kinase substrates. *C. difficile* is a Gram-positive, spore-forming, anaerobic bacterium and is the leading cause of antibiotic-associated nosocomial diarrhea (Rupnik et al., 2009). The incidence and severity of *C. difficile* infection have recently increased, representing a challenging threat to human health (Smits et al., 2016; Wiegand et al., 2012). We previously phenotypically characterized the *C. difficile prkC* mutant and highlighted the role of PrkC in CW homeostasis. A Δ*prkC* mutant exhibited modifications in cell morphology and septum formation and was also more sensitive to antimicrobial compounds that target the CW (Cuenot et al., 2019). In this work, we identified and characterized a PGH, CD1135 (renamed CwlA, see below). We showed that CwlA functions as an endopeptidase and hydrolyses PG between daughter cell to allow cell separation. We found that CwlA is phosphorylated by PrkC and demonstrated that PrkC-dependent phosphorylation controls the export of CwlA required for cytokinesis. This represents a novel and original mechanism for CW hydrolysis regulation by STK phosphorylation.

## Results

### CD1135 (CwlA) is a γ-D-Glu-mDAP-endopeptidase

CD1135 contains a signal sequence, three putative SH3_3 (also named SH3b) domains, which are predicted to contribute to PG recognition and binding (Kamitori & Yoshida, 2015; Whisstock & Lesk, 1999), and a catalytic domain of the NlpC/P60 family (**Fig. 1b**). Proteins containing NlpC/P60 domains are ubiquitous papain-like cysteine peptidases hydrolyzing the NAM-L-alanine or D-γ-glutamyl-meso-diaminopimelate linkages (Anantharaman & Aravind, 2003). Three conserved residues are involved in catalysis: Cys, His and a polar residue (His, Asp or Gln). The alignment of the NlpC/P60 domain of CD1135 with the endopeptidases LytF, LytE, CwlS, YkfC from *B. subtilis* and YkfC from *B. cereus* revealed that these residues are fully conserved (Supplementary Fig. 1). To provide evidence that CD1135 functions as a PGH, the hydrolytic activity of His_6_-tagged CD1135 protein was examined by zymogram with purified *C. difficile* PG. A clear lytic band with a molecular weight around 46 kDa corresponding to His_6_-CD1135 was detected (**Fig. 1c**), confirming the PG-degrading activity of CD1135. Many hydrolases efficiently degrade PG by acting on cross-linked PG. To characterize more precisely the PG-degrading activity and specificity of the CD1135 enzyme, we incubated the purified protein with PG of *C. difficile* 630Δ*erm*. After incubation, no soluble PG fragment were detected by reverse-phase high-pressure liquid chromatography (RP-HPLC) analysis (data not shown) and the insoluble PG fraction was further digested with mutanolysin to reveal potential cleavages inside the cross-linked PG. The resulting muropeptide profile obtained by RP-HPLC was clearly distinct from the control muropeptide profile of PG digested with mutanolysin alone (**Fig. 1d**). In particular, PG incubated with CD1135 and mutanolysin contained a major muropeptide (peak 6) present only in low amount in the PG digested with mutanolysin alone. Mass spectrometry (MS) analysis showed that peak 6 contained deacetylated disaccharide-dipeptide (Table 1), thus revealing cleavage by CD1135 of the chemical bond between g-D-Glu and mDAP inside the PG stem peptides. Concomitantly, a decrease of several muropeptides was observed in PG incubated with CD1135 compared to the control (**Fig 1d**), including monomers identified as deacetylated disaccharide-tetrapeptides with D-Ala (peak 5) or Gly (peak 2) in position 4 of the stem peptide by MS analysis (Table 1). All these results indicate that CD1135 is able to hydrolyze stem peptides inside the cross-linked PG and that CD1135 is a g-D-Glu-mDAP-endopeptidase. We decide to rename this enzyme CwlA.

**Table 1.** Proposed structures deduced from m/z values of muropeptides resulting from the digestion of *C. difficile* PG by purified His_6_-CD1135 (CwlA) followed by mutanolysin.

| Peak<sup>1</sup> | Mucopeptide<sup>2</sup> | Expected | Measured | Charge |
| --- | --- | --- | --- | --- |
|  |  | m/z | m/z | z |
| 1 | Tri (deAc) | 415.1878 | 415.1876 | 2 |
| 2 | Tri-Gly (deAc) | 443.6985 | 443.6977 | 2 |
| 3 | Tri-Ala-mDAP (deAc) | 536.7487 | 536.7463 | 2 |
| 4 | Di | 699.2936 | 699.2934 | 1 |
| 5 | Tetra (deAc) | 450.7064 | 450.7062 | 2 |
| 6 | Di (deAc) | 657.2831 | 657.2833 | 1 |
| 7 | Tri-Tri-Gly (deAc X2) | 848.8733 | 848.8749 | 2 |
| 8 | Tri-Tetra (deAc X2) | 855.8811 | 855.8809 | 2 |
| 9 | Tri-Tetra (deAc X2) | 855.8811 | 855.8823 | 2 |
| 10 | Tetra-Tetra (deAc X2) | 891.3997 | 891.4020 | 2 |
| 11a | Tetra-Tri-Tri (deAc X3) | 841.0398 | 841.0406 | 3 |
| 11b | Tetra-Tetra (deAc X2) | 891.3997 | 891.4010 | 2 |
| 12 | Tri-Tetra-Tetra (deAc X3) | 864.7189 | 864.7201 | 3 |

<sup>1</sup>Peak number correspond to Fig. 1d
<sup>2</sup>Di, disaccharide dipeptide (L-Ala-D-Glu); Tri, disaccharide tripeptide (L-Ala-D-iGlu-mDAP); Tetra, disaccharide tetrapeptide (L-Ala-D-iGlu-mDAP-D-Ala); disaccharide, GlcNAc-MurNAc; deAc, deacetylation on GlcNAc, iGlu, isoglutamic acid; mDAP, mesodiaminopimelic acid, Gly, glycine.

### Changes in CwlA abundance leads to cell separation defects during growth

To determine the role of this PGH in *C. difficile*, a *cwlA* mutant was generated. This mutant grew similarly to the 630Δ*erm* wild-type (WT) strain (Supplementary Fig. 2a). Using phasecontrast microscopy, we revealed during the exponential growth a slightly but significant increase in cell length for the mutant (7.8 ±2.4 μm) as compared to WT cells (6.5 ±1.6 μm) (**Fig. 2a,b**). 50% of the mutant cells also existed as unseparated paired cells compared with only 15% for the WT strain (**Fig. 2a,c**). Cell length and cell separation were almost completely restored in a *cwlA* mutant complemented with a plasmid harboring the *cwlA* gene expressed from its own promoter (**Fig. 2a-c**). This result strongly suggests that the PGH, CwlA, is required to cleave PG at the septum to allow separation of daughter cells. To determine the localization of CwlA, we constructed a plasmid encoding a CwlA-SNAP^*Cd*^ fusion produced under the control of the anhydrotetracycline (ATc)-inducible P_*tet*_ promoter. After 2 h of induction in the presence of 50 ng/ml of ATc, the CwlA-SNAP^*Cd*^ fusion protein was distributed through the cytoplasm but enriched at the septum of dividing cells (**Fig. 2d**). However, transmission electron microscopy (TEM) analysis of WT, *cwlA* mutant and complemented cells revealed no difference in septa and CW thickness (Supplementary Fig. 2b). These results indicate that CwlA is involved in cytokinesis.

**Figure 2.**
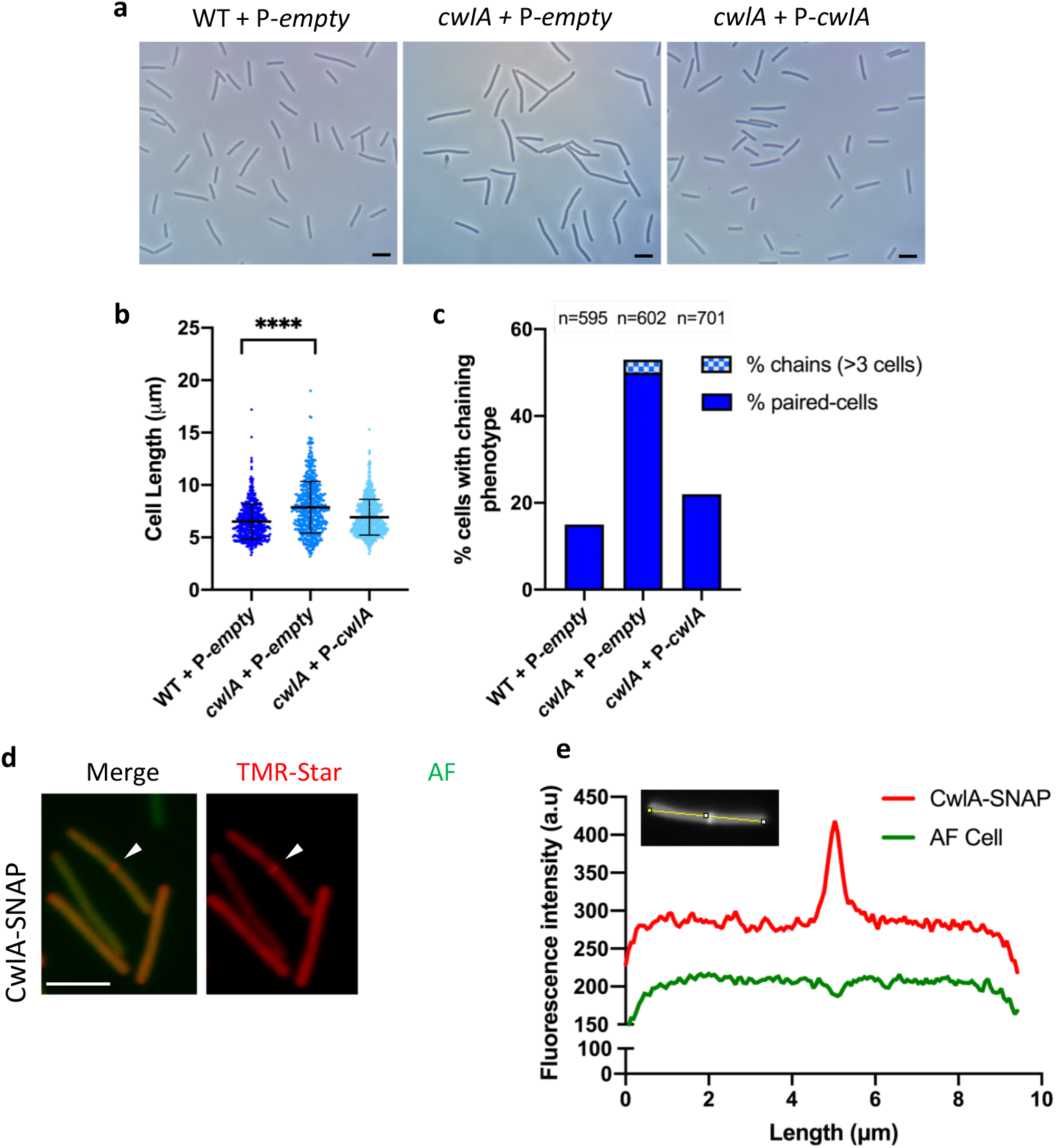
CwlA is involved in cell division. **a**, Phase contrast images of exponentially growing *C. difficile* cells of strains 630Δ*erm* (WT) + P-*empty, cwlA* + P-*empty, cwlA* + *P-cwlA*. Scare bar, 5 μm. **b**, Scatter plots showing cell length with the median and standard deviation (SD) of each distribution indicated by a black line. *P* value was determined by two-sided Mann–Whitney *U* tests (\*\*\*\**P* < 0.0001); counted 575 (WT + P-*empty*), 602 (*cwlA* + P-*empty*), 701 (*cwlA* + P-*cwlA*). **c**, Percentage of cells harboring a chaining phenotype, the n value represents the number of cells analyzed in a single representative experiment. The images are representative of experiments performed in triplicate. **d**, CwlA-SNAP protein fusion and its localization. A plasmid carrying the SNAP-fusion was introduced into the *CwlA* strain. Merge images (*left row*), TMR-Star fluorescent signal (*middle row*) and auto-fluorescence AF (*right row*). Scale bar, 5 μm. **e**, Quantification of fluorescence in arbitrary units (a.u) along the cell for a typical cell.

We then expressed *cwlA* under the control of the P_*tet*_ inducible promoter on a plasmid. Growth of the *cwlA* mutant containing pDIA6103-P_*tet*_ *cwlA* was impaired in the presence ATc (**Fig. 3a, b**). In addition, we observed a filamentation phenotype upon induction of *cwlA* expression with 50 ng/ml of ATc (**Fig. 3c** and Supplementary Fig. 3). By contrast, neither WT nor *cwlA* mutant strains harboring an empty plasmid were similarly affected in morphology or in growth (**Fig. 3b,c** and Supplementary Fig. 3). The filamentation phenotype in strains that overexpress a gene encoding an endopeptidase is unusual (Uehara & Bernhardt, 2011). Staining with FM4-64 and TEM revealed few septa in these filamented cells, suggesting that division was impaired prior to constriction (**Fig. 3c,d**). We then compared the PG structure of the WT strain, the *cwlA* mutant and cells overexpressing *cwlA*. Purified PG was digested with mutanolysin to generate muropeptides for RP-HPLC analysis. However, no significant differences were detected between these strains, suggesting that the phenotype observed in cells overexpressing *cwlA* is not the result of a higher CwlA activity (Supplementary Fig. 4). The observed filamentation might be therefore due to an impaired access to PG of other PGH containing SH3_3 domains by protein-protein interaction via their SH3_3 domains. To test this possibility, we expressed under the control of the P_*tet*_ promoter the 5’ part of the *cwlA* gene corresponding to the three SH3_3 domains associated to the signal peptide. The *cwlA* mutant overexpressing this truncated gene also exhibited elongation and growth defect phenotypes (**Fig. 3b,c** and Supplementary Fig. 3). SH3_3 domains are thus responsible for the cell division defect observed upon *cwlA* overexpression.

**Figure 3.**
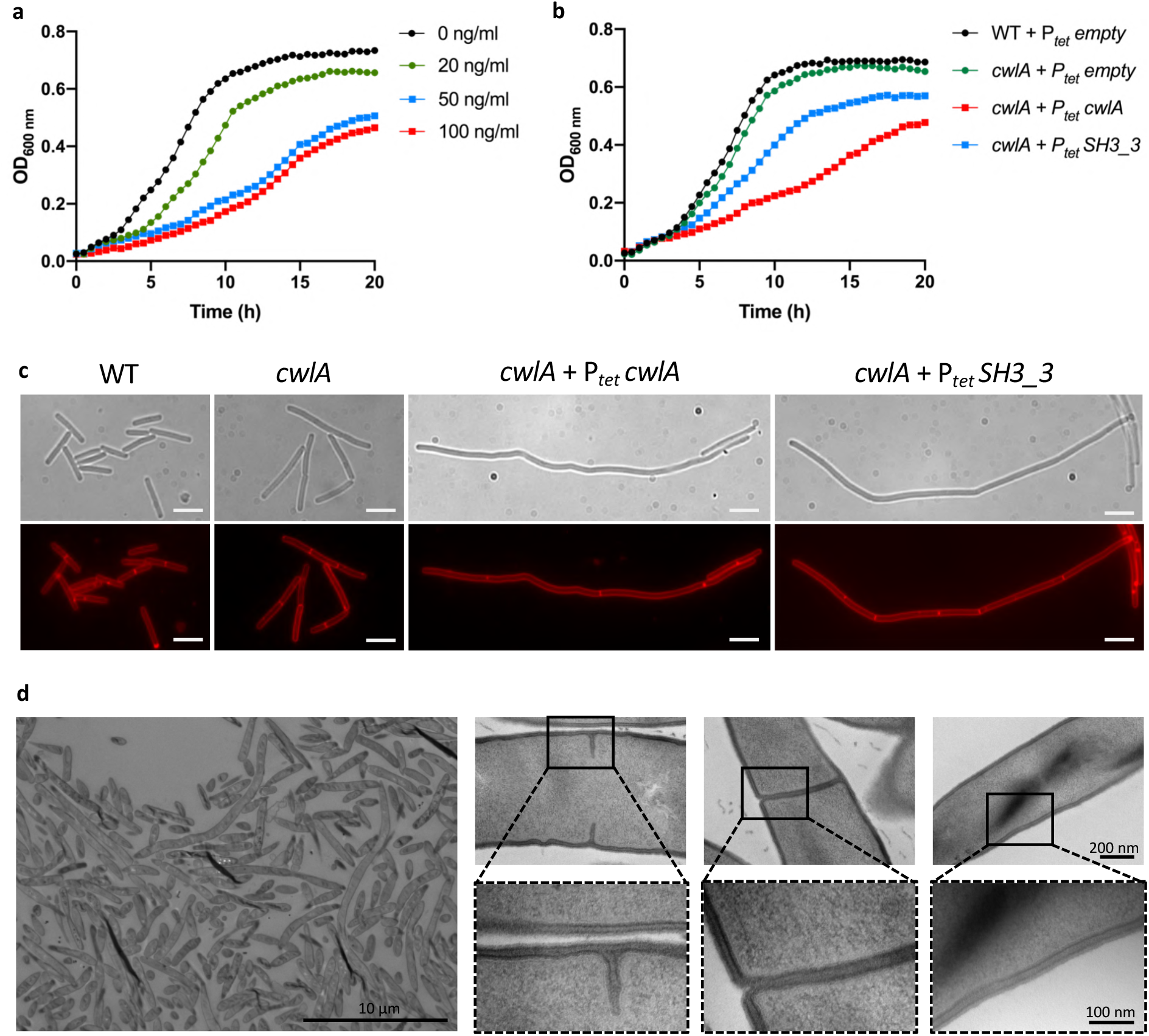
Overexpression of *cwlA* inhibits growth and produces filamented cells. **a,** Growth curves of *cwlA* mutant carrying pDIA6103-P_*tet*_ *cwlA* in TY at 0, 20, 50 and 100 ng/ml ATc. **b**, Growth curves showing the effect of overexpression of *cwlA* or only its SH3_3 domains when 50 ng/ml of ATc was added at the beginning of growth. **c**, Phase contrast images (*upper row*) and fluorescence microscopy (*lower row*) of cells overexpressing *cwlA* full length or only the *SH3_3* domains stained with FM4-64. Scale bars, 5 μm. **d,** Transmission electron micrographs of septal regions and cell wall from cells overexpressing *cwlA*. Lower panels: higher magnifications (100 nm) of the division septa or cell wall highlighted by a black square. Growth curves and images are representative of experiments performed in triplicate.

### *CD2148* and *prkC* mutants show similar phenotypes to lack and excess of CwlA, respectively

The *C. difficile* genome contains genes encoding two STKs, the PASTA-kinase PrkC and a second kinase, CD2148, and the associated PP2C-type phosphatase (STP) (**Fig. 4a**). We previously obtained a *prkC* deletion mutant (Cuenot et al., 2019). We constructed here a *stp* deletion as well as a *CD2148* mutant in both 630Δ*erm* and Δ*prkC* strains. Interestingly, *prkC* inactivation led to cell filamentation (Cuenot et al., 2019) (**Fig. 4b,c**), a phenotype reminiscent to bacteria overexpressing *cwlA* (**Fig. 3**). In contrast, a *CD2148* mutant exhibited a slight increase in cell length (10.5 ± 4.0 μm) compared to the WT strain, similarly to the *cwlA* mutant, and a cell separation defect more accentuated than in the *cwlA* mutant (**Fig. 4b,c** and Supplementary Fig. 5). In the *CD2148* mutant, 98% of the cells presented a chaining phenotype with 53% of cells found as chains of more than 3 unseparated cells **(Fig. 4c, right**). The WT phenotype was recovered upon introduction of a plasmid containing the *CD2148* gene under the control of its own promoter (Supplementary Fig. 5c,d). The *stp* deletion mutant showed a filamentation phenotype with an average size corresponding to 13.7 ± 6.6 μm and 37% of cells were found as paired-cells (**Fig. 4b,c**).

**Figure 4.**
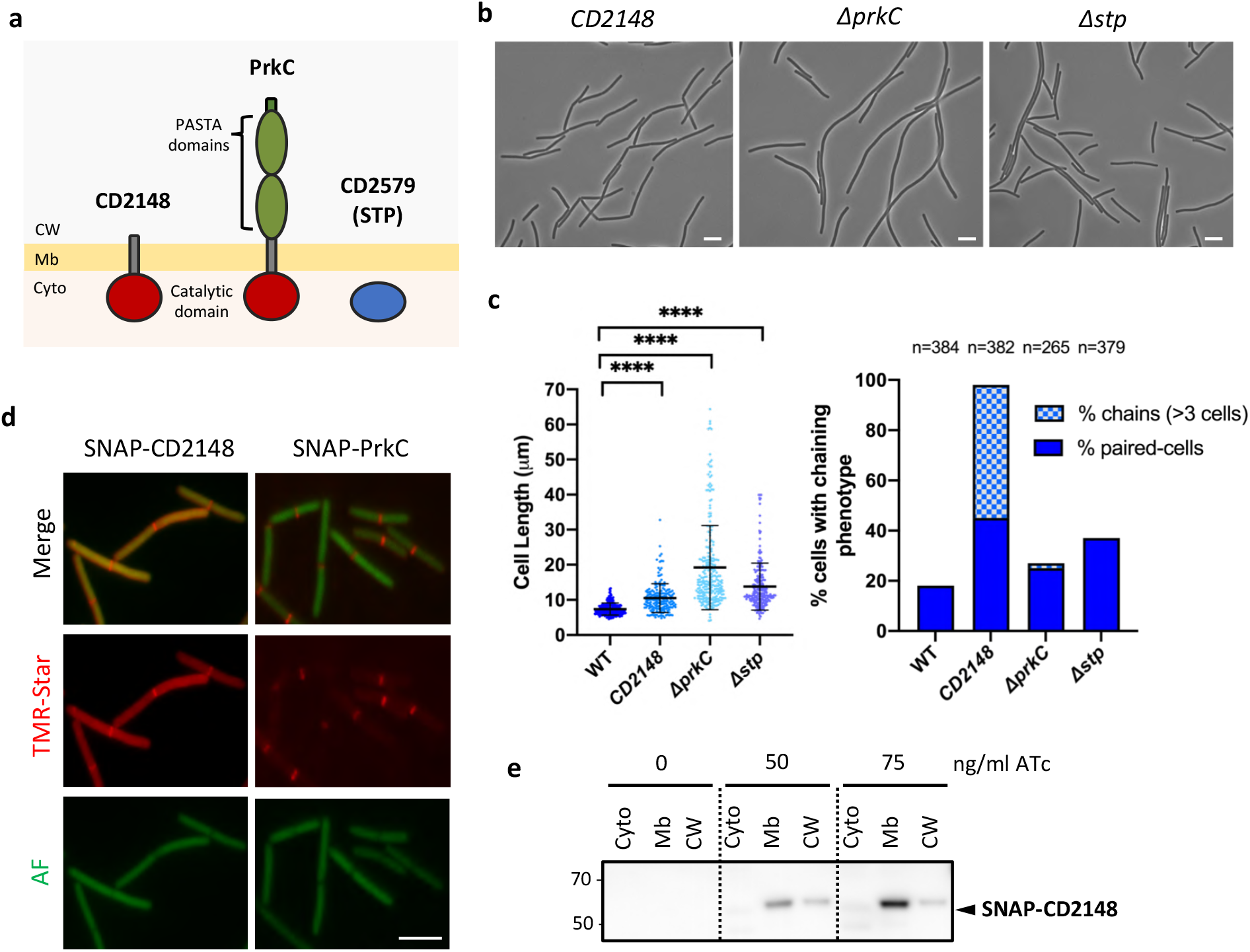
*CD2148* and Δ*prkC* mutants show similar phenotypes to lack or overproduction of CwlA. **a,** Schematic representation of *C. difficile* STKs. **b**, Phase contrast images of *CD2148*, Δ*prkC* and Δ*stp* mutants. Scare bar, 5 μm. **c**, Scatter plots showing cell length of STK mutants (*left panel*) and percentage of cells harboring a chaining phenotype (*right panel*). *P* values were determined by two-sided Mann-Whitney *U* tests (\*\*\*\**P* < 0.0001); counted 320 (WT), 188 (*CD2148*), 251 (Δ*prkC*) and 231 (Δ*stp*). **d**, Localization of SNAP-CD2148 and SNAP-PrkC protein fusions. Merge images (*upper row*), TMR-Star fluorescent signal (*middle row}* and auto-fluorescence AF (*lower row*). Scale bar, 5 μm. The images are representative of experiments performed in triplicate. **e**. Western blot showing the presence of SNAP-CD2148 in the different cell fractions: cytoplasm (Cyto), membrane (Mb) and cell wall (CW).

Localization of CD2148 was then determined using a plasmid carrying a P_*tet*_ SNAP^*Cd*^-*CD2148* fusion. After induction, we detected the SNAP-CD2148 fusion protein in the cytoplasm with an enrichment at mid-cell (**Fig. 4d**). This localization pattern is similar to that of CwlA (**Fig. 2d**) but different from that of PrkC, which mostly localized at the cell septum (**Fig. 4d**). These results indicated that CD2148, similarly to CwlA, partially localizes at the cell septum in agreement with its role in cell separation. In contrast to PrkC, the CD2148 kinase does not contain an extracellular PASTA domain but the presence of a transmembrane segment at the C-ter of the protein suggests a localization at the membrane (**Fig. 4a**) (Cuenot et al., 2019; Madec et al., 2003; Morlot et al., 2013). To analyze the cellular localization of CD2148, we also performed a cell fractionation to separate cytoplasmic, membrane and CW fractions.

Western blots using antibodies raised against SNAP revealed that CD2148 is mainly found in the membrane (**Fig. 4e**).

### CwlA is phosphorylated *in vivo* and *in vitro* by PrkC

Since PrkC and CD2148 are STKs, we therefore compared the level of phosphorylation of CwlA in the WT, Δ*prkC*, *CD2148*, Δ*prkC CD2148* and Δ*stp* strains. CwlA was phosphorylated at S136 in all strains. However, a higher intensity of S136 phosphorylation was detected in the Δ*stp, CD2148* and double mutants (**Fig. 5a, left panel**). A second residue, the T405, was also found phosphorylated in the WT strain as well as in the Δ*stp* and *CD2148* mutants while its phosphorylation was abolished in the Δ*prkC* mutant and in the double mutant (**Fig. 5b, right panel**). These results indicated that T405 phosphorylation is PrkC-dependent *in vivo*. Furthermore, we observed that the level of phosphorylation of T405 was significantly higher (FC>2 in log2) in the Δ*stp* and *CD2148* mutants compared to the WT strain while the peptide amounts were similar. STP is probably involved in the desphosphorylation of CwlA. In addition, the increased phosphorylation level of T405 detected in the *CD2148* mutant disappeared in the double *CD2148* Δ*prkC* mutant. This result suggested that the effect of *CD2148* inactivation on T405 phosphorylation was mediated through PrkC and that the role of the second STK, CD2148, was more complex than expected (see below). Moreover, additional phosphorylated residues were detected in the *CD2148* and *Δstp* mutants (Supplementary Fig. 6a). This may contribute to an hyperphosphorylation of CwlA in these strains. To confirm the specificity of CwlA phosphorylation *in vitro*, we purified the catalytic domain of PrkC and CD2148. Purified CwlA was incubated with the purified kinase domain of PrkC or CD2148 at a 1:10 ratio in presence of cold ATP. To determine the phosphorylation status of CwlA (**Fig. 5c**), a Phos-Tag™ fluorescent gel stain method allowing the specific detection of phosphorylated proteins was used (Kusamoto et al., 2019). When CwlA was incubated with PrkC, two bands with a close molecular weight were detected. The upper band corresponded to CwlA (46.1 kDa) and the lower band to the kinase domain of PrkC (45 kDa), which is known to be autophosphorylated (Madec et al., 2003; Nováková et al., 2005; Young et al., 2003). By contrast, no fluorescence was detected when CwlA was incubated alone or with CD2148 (**Fig. 5c**). To detect phosphorylated isoforms of purified His_6_-CwlA, we further used Phos-Tag™ Acrylamide gel to separate phosphorylated and non-phosphorylated forms (Kinoshita et al., 2006). PrkC was the only kinase that efficiently phosphorylated CwlA *in vitro* (**Fig. 5d** and Supplementary Fig. 6b). The phosphorylation of S136 and T405 detected *in vivo* was then confirmed *in vitro* by mass spectrometry (**Fig 5e**). T405, located within the NlpC/P60 domain, was more specific for PrkC-dependent phosphorylation in agreement with our *in vivo* results where this phosphorylation was found strictly PrkC-dependent. In order to confirm whether T405 was specifically phosphorylated by PrkC, this residue was replaced with the non-phosphorylable residue alanine (T405A) or the phospho-mimetic residue aspartate (T405D) by site directed mutagenesis. The CwlA-T405A and CwlA-T405D proteins were then purified and tested *in vitro* for PrkC-mediated phosphorylation. In both cases, *in vitro* phosphorylation of CwlA-T405A and CwlA-T405D by PrkC was abolished, as determined by Phos-Tag™ Acrylamide (Supplementary Fig. 6b). Altogether these results indicated that T405 is the major site of STK-dependent phosphorylation of CwlA and that this phosphorylation is specific of PrkC.

**Figure 5.**
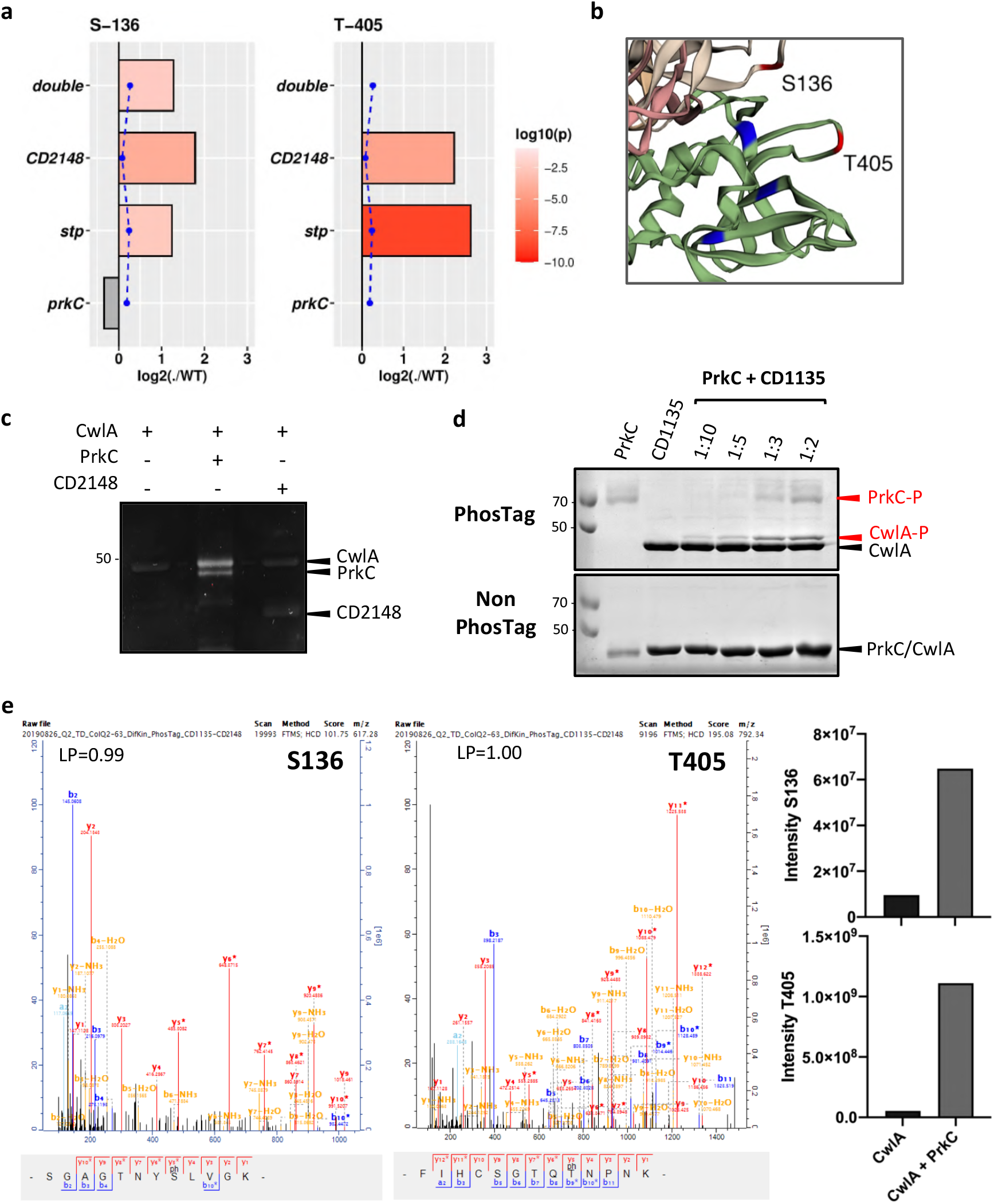
CwlA is phosphorylated by PrkC. **a,** *In vivo* phosphorylation of CwlA as detected in phosphosproteomic data. Bar graphs represent fold change of the phosphosite Serine 136 (S-136) or Threonine 405 (T-405) among Δ*prkC*, Δ*stp, CD2148* and Δ*prkC CD2148* (*double*) mutants as compared to WT. An absence of bars means that no phosphopeptides were detected. Significant and not significant are represented by log10 (P-value) in red or gray color, respectively. Blue dotted line represents fold change of peptide amounts. **b**, Phosphorylated residue S136 and T405 (in red) are in close proximity with the three conserved catalytic residues: Cys, His and Asp (in blue). **c**, *In vitro* phosphorylation assay of CwlA by the purified STKs, PrkC and CD2148. Phosphorylation was visualized by Phos-tag™ fluorescent gel stain reagents. **d**, *In vitro* phosphorylation assay of CwlA by PrkC at different ratio of kinase:substrate. Phosphorylation was visualized by Phos-tag™ Acrylamide for the separation of phosphorylated and nonphosphorylated proteins. **e**, Mass spectrometry detection of CwlA phosphorylation sites after *in vitro* phosphorylation reactions performed without or with PrkC. LP is the localization probability of the identified phosphosite.

### Phosphorylation at T405 does not impact CwlA PGH activity

Based on the 3D structure prediction of CwlA obtained with Phyre 2.0 (Kelley et al., 2015), the phosphorylated residues S136 and T405 are likely located in the second SH3_3 and in the NlpC/P60 domain, respectively (Supplementary Fig. 6a). T405 is in the vicinity of the three conserved catalytic residues, suggesting that PrkC-dependent phosphorylation could regulate CwlA activity (**Fig 5b** and Supplementary Fig. 1b). To determine if the phosphorylation could modify the PG-degrading activity, the purified CwlA-T405A and CwlA-T405D proteins or the CwlA protein phosphorylated *in vitro* by PrkC were tested by zymogram. Neither the substitution of T405 nor the phosphorylation of CwlA by PrkC affected the hydrolytic activity of these proteins in zymogram (**Fig. 1c**). These results indicate that T405 phosphorylation is neither beneficial nor detrimental for the activity of CwlA.

### Effect of T405 phosphomimetic and non-phosphorylable mutations of CwlA *in vivo*

To study the role of the T405 phosphorylation, we expressed in the *cwlA* mutant, genes encoding CwlA, CwlA-T405A or CwlA-T405D fused to a hemagglutinin (HA) tag under the control of the inducible P_*tet*_ promoter. Cells were grown for 3 h before adding 50 ng/ml of ATc inducer for 2 h. Cell length and separation defect were restored in cells expressing a WT copy of *cwlA* (**Fig. 6a-c** and Supplementary Fig. 7) or *cwlA-T405A*. In contrast, cells expressing *cwlA-T405D* retained a defect in cell length and in cell separation, similarly to cells carrying an empty plasmid (**Fig. 6a-c**). To localize CwlA-HA in the different cellular fractions and determine the impact of these mutations, we analyzed by western blot the distribution of CwlA in the different cell fractions (cytoplasm, membrane and CW). As a fractionation control, we used antibodies raised against Cwp66, a CW-associated protein of *C. difficile* (Fagan & Fairweather, 2014; Karjalainen et al., 2001; Waligora et al., 2001). Cwp66 was mainly detected in the CW fraction (**Fig. 6d**) as expected. Using an anti-HA antibody, we detected similar amount of CwlA-HA in the cytoplasm and in the membrane for the three strains. In contrast, a significantly reduced amount of CwlA-T405D and a slightly increased amount of CwlA-T405A were detected in the CW compared to the CwlA protein (**Fig. 6d**). These results suggest that phosphorylation plays a role in controlling the export of the CwlA PGH.

**Figure 6.**
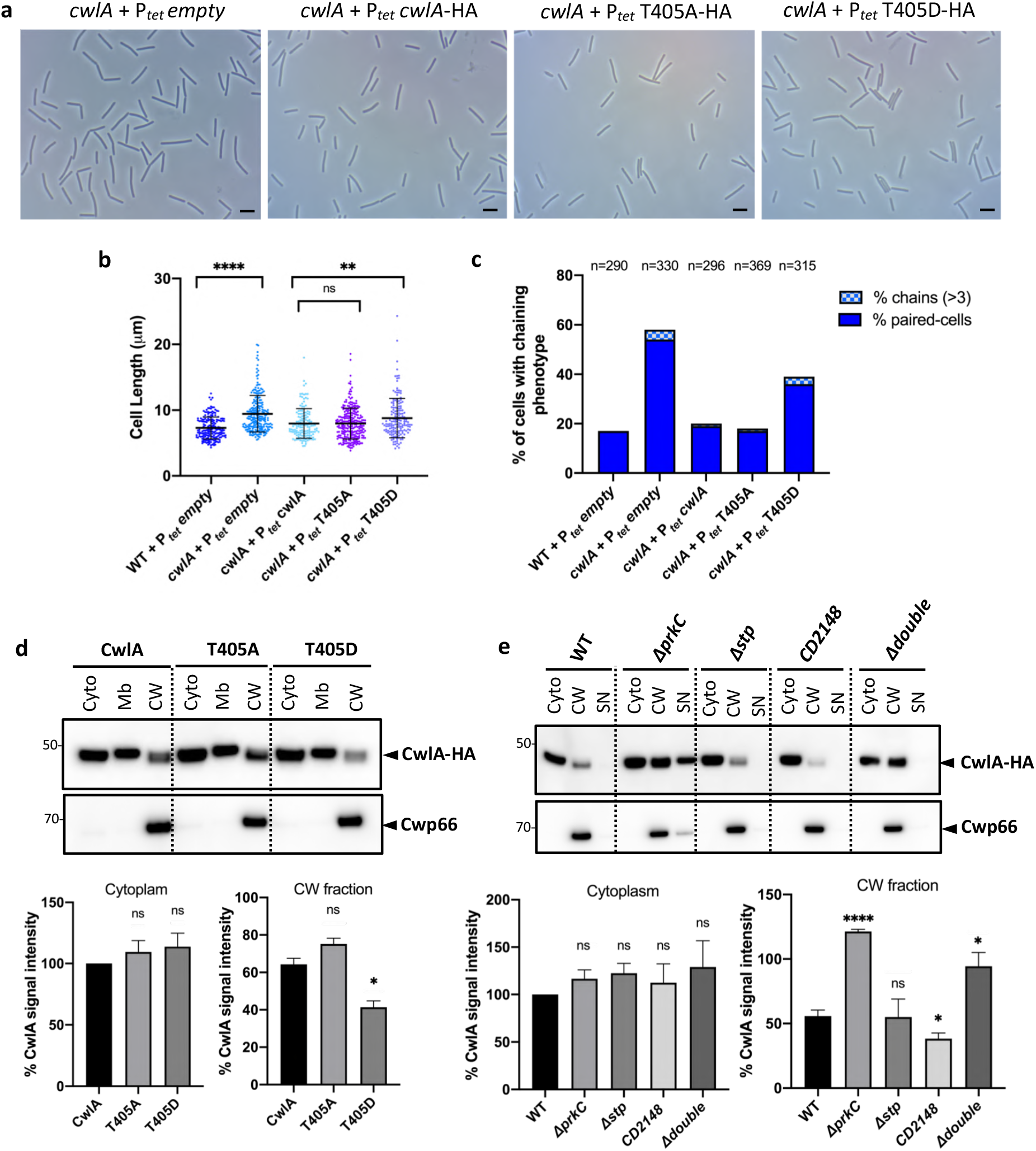
Effect of CwlA phosphorylation on its cell wall localization and on cell division. **a,** Phase contrast images of *C. difficile* cells of strains *cwlA* + P_*tet*_ *empty, cwlA* + P_*tet*_ *cwlA*-HA, *cwlA* + P_*tet*_ *cwlA-T405A*-HA and *cwlA* + P_*tet*_ *cwlA-T405D*-HA. Scale bar, 5 μm. **b**, Scatter plots showing the distribution of cell length for each strain. Two-sided Mann-Whitney *U* tests (\*\*\*\**P* < 0.0001; ** P< 0.01; ns = 0.9143), cells counted: 190 (WT + P_*tet*_ *empty*), 230 (*cwlA* + P_*tet*_ *empty*), 146 (*cwlA* + P_*tet*_ *cwlA*-HA), 269 (*cwlA* + P_*tet*_ *T405A*-HA) and 167 (*cwlA* + P_*tet*_ *T405D*-HA). **c**, Percentage of cells harboring a chaining phenotype, n indicates the number of cells counted per strain in a single representative experiment. **d**, (*upper panel*) Western blots showing the levels of CwlA, CwlA-T405A and CwlA-T405D proteins in the different cell fractions: cytoplasm (Cyto), membrane (Mb) and cell wall (CW). Cwp66 was used as a fractionation control. These proteins were detected using anti HA and anti Cwp66 antibodies, respectively. (*lower panel*) Quantification of CD1135 signals by densitometry. The CwlA value of cytoplasm fraction was arbitrarily set to 100 and asterisks indicate statistically significant differences. From left to right (Cytoplasm): *P* = 0.291 and 0.221 (ns). From left to right (Cell wall fraction): *P* = 0.075 (ns) and 0.020 (\**P* < 0.05). **e**, (*upper panel*) Western blots showing the levels of CwlA-HA in the different cell fractions of STK and STP mutants: cytoplasm (Cyto), cell wall (CW) and supernatants (SN). (*Lower panel*) Quantification of signals by densitometry. From left to right (Cytoplasm): *P* = 0.126, 0.070, 0.478 and 0.265 (ns). From left to right (Cell wall fraction): *P* = 0.000011 (\*\*\*\**P* < 0.0001), 0.275 (ns), 0.014 and 0.44 (\**P* < 0.05). Average values and standard error of the means are calculated from 2 independent experiments in d and from 4 independent experiments in e.

### PrkC and CD2148 influence the abundance of cell wall-associated CwlA

To test the influence of phosphorylation on the export of CwlA, we investigated the cellular location of CwlA-HA in strains expressing P_*tet*_ *cwlA*-HA and inactivated for the STKs or STP. We detected CwlA-HA in both the cytoplasm and the CW in all the tested strains. While the amount of CwlA-HA detected in the cytoplasm was stable, we observed variations of its abundance in the CW fraction (**Fig. 6e**). CwlA-HA levels increased approximately 2-fold in the CW of the Δ*prkC* and the double Δ*prkC CD2148* mutants compared to the WT strain. By contrast, the amount of CwlA-HA was significantly reduced in the CW of the *CD2148* mutant. Levels of CwlA-HA in the CW of the *stp* mutant were similar to those of the WT strain (**Fig. 6e**). Because reduced CwlA levels were found in the CW of *CD2148* mutant, we tested whether CwlA was present in the supernatant fraction. However, CwlA was shed into the supernatant only for the Δ*prkC* mutant. This is probably due to steric hindrance caused by the high abundance of CwlA attached to the CW and the increased lysis of Δ*prkC* cells (Cuenot et al., 2019). Altogether, these results supported that the ability of CwlA to cross the membrane, and thus exert PG hydrolysis on the CW is dependent on its phosphorylation level.

### Overexpression of *cwlA* restores the cell separation defect in *CD2148* cells

Cell division defects observed in the *CD2148* mutant are probably due to the decreased level of CwlA in the CW (**Fig. 6e**). We next investigated whether *cwlA* overexpression in the *CD2148* mutant could restore a WT phenotype. The *CD2148* mutant carrying the P_*tet*_ *cwlA*-HA on a plasmid was grown for 3 h before inducing the expression of *cwlA* with a range of ATc concentrations. In absence of induction, 97% of the cells exhibited a chaining phenotype, 55% with 3 or more unseparated cells. After induction with 20 ng/ml of ATc, 83% of cells exhibited separation defects with 27% as chains of at least 3 cells. However, in presence of 50 or 100 ng/ml of ATc, we observed a restoration of cell length and 41 or 39% of paired-cells respectively and only 2% as chains of more than 3 cells (**Fig. 7a,b**). To correlate these results to the accumulation of CwlA in the CW, CwlA levels were determined by Western blotting. In presence of 20 ng/ml of ATc, CwlA was barely detected in the CW, in agreement with the phenotype observed in this condition. However, the level of CwlA in the CW increased substantially when CwlA expression was induced with 50 or 100 ng/ml ATc (**Fig. 7c**), these amounts being sufficient to partially restore a WT phenotype. Overexpression of *cwlA* compensates for the absence of CD2148. This might be due to an increase of the pool of nonphosphorylated CwlA present in the CW restoring cell separation.

**Figure 7.**
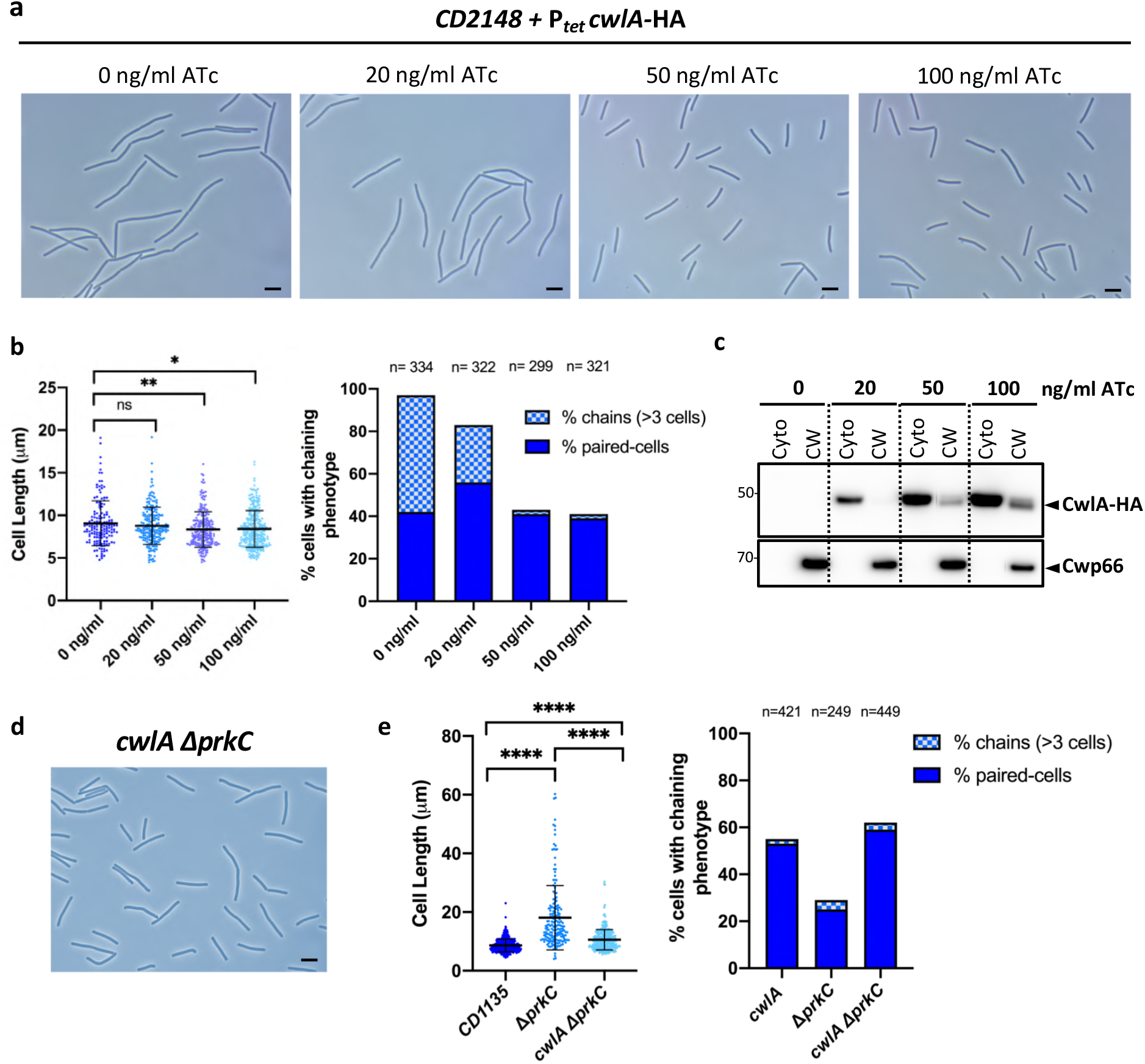
PrkC and CD2148 controls export of CwlA to the cell wall. **a,** Phase contrast images of exponentially growing *CD2148* cells expressing *cwlA*-HA under the control of a P_*tet*_ promoter at different ATc concentrations (0, 20, 50 and 100 ng/ml). Scale bar, 5 μm. **b**, Scatter plots showing cell length with the median and SD of each distribution indicated by a black line. *P* values were determined by two-sided Mann-Whitney *U* tests. *P*= 0.7146 (ns), 0.0074 (\*\**P* < 0.01) and 0.0116 (\**P* < 0.05); counted 152 (0 ng/ml), 197 (20 ng/ml), 236 (50 ng/ml) and 287 (100 ng/ml) (*left panel*). Percentage of cells harboring a chaining phenotype, n indicates the number of cells counted per strain in a single representative experiment (*right panel*). **c**, Western blots showing the levels of CwlA-HA in the cytoplasm (Cyto) and cell wall (CW) of *CD2148* mutant at increasing concentrations of ATc. **d**, Phase contrast images of the double Δ*prkC cwlA* mutant. Scale bar, 5 μm. **e**, Scatter plots showing cell length (*left panel*) and percentage of cells harboring a chaining phenotype (*right panel*). Two-sided Mann-Whitney *U* tests (\*\*\*\**P* < 0.0001; counted 431 (*cwlA*), 183 (Δ*prkC*) and 329 (Δ*prkC cwlA*).

### Cell elongation is restored in the Δ*prkC cwlA* double mutant

The cell division and morphology defects observed in the Δ*prkC* mutant could also be linked to the level of the PGH, CwlA, in the CW. To determine if the high level of CwlA in the CW of the Δ*prkC* mutant triggers cell elongation, we constructed a double *cwlA* Δ*prkC* mutant. Phase-contrast microscopy analysis of *cwlA* Δ*prkC* mutant cells revealed a significant reduction in cell length (10.5 ± 3.4 μm) compared to Δ*prkC* cells (18.1 ± 10.9 μm) and a slight but significant increase in size compared to the *cwlA* mutant (8.6 ± 2.1 μm) (**Fig. 7d,e**). The double *cwlA* Δ*prkC* mutant also presented a chaining phenotype similar to the *cwlA* mutant (**Fig. 7e, right**). These results indicate that elongation of cells in the Δ*prkC* mutant is mainly due to the accumulation of CwlA in the CW. This accumulation was also responsible for the increased susceptibility of Δ*prkC* mutant to antimicrobial compounds targeting CW including two cephalosporins (cefoxitine and ceftazidime), teicoplanin and bacitracin (Table 2).

**Table 2.** MICs of the mutants used in this study for antibiotics targeting CW

| Antibiotics | MIC (μg/ml) |  |  |  |  |  |
| --- | --- | --- | --- | --- | --- | --- |
|  | 630 *Δerm* | *ΔprkC* | *Δstp* | *CD2148* | *cwlA* | *cwlA ΔprkC* |
| <u>Cephalosporins</u> |  |  |  |  |  |  |
| Cefoxitin | >256 | 48 | >256 | >256 | >256 | >256 |
| Ceftazidime | 64 | 3 | 64 | 96 | 96 | 64 |
| <u>Other Cell Wall</u> |  |  |  |  |  |  |
| Teicoplanin | 0,12 | <0,016 | - | - | 0,12 | 0,12 |
| Bacitracin | >256 | 8 | >256 | >256 | >256 | >256 |

## Discussion

In this study we characterized a new PGH of *C. difficile*, CwlA, that cleaves the PG and plays a role in cell division. In addition, we identified the STK PrkC as a direct regulator for this hydrolase by modulating CwlA export and access to the CW, highlighting a novel regulatory mechanism to control CW hydrolysis by phosphorylation.

When bacteria divide, they form a septum and each daughter cell contains a membrane with a layer of PG that is shared between them. To separate the cells and complete the cell division process, the shared PG must be partially hydrolyzed by dedicated enzymes (Adams & Errington, 2009). CwlA belongs to the NlpC/P60 family, a major class of CW degrading enzymes that typically function as endopeptidases and hydrolyze γ-D-glutamyl-*meso*-diaminopimelate or NAM-L-alanine linkages (Anantharaman & Aravind, 2003). Here, we demonstrated that CwlA cleaves the cross-linked PG at the level of stem peptides. We identified the first PGH involved in cell division in *C. difficile* allowing separation of daughter cells. Accordingly, around 50% of cells were found as a chain of paired cells in a mutant lacking *cwlA*. Even if phenotypes are sometimes difficult to observe with a single PG hydrolase gene knockout probably due to functional redundancy (Vollmer, Joris, et al., 2008), mutants lacking splitting hydrolases can fail to properly divide and form chains of unseparated cells as observed for the *cwlA* mutant. An *atl* mutant of *S. aureus* and a *lytB* mutant of *S. pneumoniae* form clusters of non-separated cells. These hydrolases are localized at the sites of cell division (García et al., 1999; Takahashi et al., 2002) as observed for CwlA. In *B. subtilis*, LytE, LytF, CwlS or CwlO hydrolyze the linkage of γ-D-glutamyl-*meso*-diaminopimelic acid in PG (Fukushima et al., 2006; Schmidt et al., 2001; Yamaguchi et al., 2004). LytE and CwlO break the PG along the lateral cell wall to support cell elongation while LytF and CwlS are implicated in cell separation (Domínguez-Cuevas et al., 2013; Fukushima et al., 2006; Yamamoto et al., 2003). Inactivation of either *cwlO* or *lytE* produces shorter cells, and a double mutant is lethal (Hashimoto et al., 2012). In contrast to LytF, LytE and CwlS, which contain several LysM PG-binding domains, CwlA possesses a less-characterized SH3 PG-binding domain (Desvaux et al., 2018). Recently, it has been shown that the unclassical SH3 domain of lysostaphin recognizes the pentaglycine cross-bridges present in staphylococcal PG, positioning the enzyme to cleave the cross-bridges (Gonzalez-Delgado et al., 2020). However, these crossbridges are absent in the PG of *C. difficile* (Peltier et al., 2011), indicating a different SH3-binding motif. Overproduction of highly active PGHs usually results in cell lysis (Uehara & Bernhardt, 2011). Surprisingly, we rather observed that overexpression of *cwlA* leads to cell elongation. The elongation phenotype that we observed is associated to the overproduction of the SH3_3 domains, which could impair the access of other PGHs to PG. Interestingly, *C. difficile* possesses three other NlpC/P60 endopeptidases associated to one to three SH3 domains (Wydau-Dematteis et al., 2018) (Supplementary Fig. 8) that may have redundant functions with CwlA. A fourth PGH, Acd, that hydrolyses peptidoglycan bonds between NAG and NAM (Dhalluin et al., 2005) is associated to four SH3 domains.

Proper PG hydrolysis is essential for cell wall biogenesis. However, due to their destructive potential, PGH activity must be tightly regulated to ensure that hydrolases act when and where they should in coordination with PG synthesis to prevent CW damages (Do et al., 2020; Uehara & Bernhardt, 2011; Vermassen et al., 2019). In this study, we uncovered a novel and original mechanism for PG hydrolysis control by STK-dependent phosphorylation. We demonstrated that CwlA is specifically phosphorylated at T405 *in vivo* by the PASTA-STK, PrkC. Further *in vitro* phosphorylation using the purified kinases PrkC and CD2148 confirmed the T405 phosphorylation by PrkC. In other Gram-positive bacteria, PASTA-STKs rather control the expression of endopeptidase genes or other genes involved in cell wall metabolism through transcriptional regulators like WalR-WalK and GraR. PrkC of *B. subtilis* phosphorylates WalR, which controls *lytE* and *cwlO* genes encoding endopeptidases in early stationary phase (Dobihal et al., 2019; Libby et al., 2015). StkP in *S. pneumoniae* has been also proposed to function in concert with the WalK through protein-protein interaction (Stamsås et al., 2017). PASTA-STKs also phosphorylate enzymes involved in PG metabolism such as Glm or Mur enzymes and the flippase MviM (Fridman et al., 2013; Gee et al., 2012; Hardt et al., 2017; Nováková et al., 2005; Parikh et al., 2009; Pensinger et al., 2018). Interestingly, the amidase CwlM is phosphorylated by the STK, PknB, in *M. tuberculosis*. However, in contrast to CwlA, which is exported to the CW, CwlM remains in the cytoplasm and its phosphorylation stimulates the catalytic activity of MurA, the first enzyme in the PG precursor synthesis pathway (Boutte et al., 2016). Finally, several proteins involved in cell division such as DivIVA, MapZ, GspB or FtsZ are also phosphorylated by STKs in *B. subtilis, S. pneumoniae* or *S. aureus* (Fleurie, Lesterlin, et al., 2014; Fleurie, Manuse, et al., 2014; Manuse et al., 2016; Pensinger et al., 2018; Pereira et al., 2011).

We evidenced that CwlA phosphorylation by PrkC does not affect its catalytic function but rather inhibits its export. Thereby, in this work, we provide a novel mechanism of control of CW homeostasis by the STKs. In our proposed model (**Fig. 8**), the homeostatic control of CwlA ensures that growing cells maintain a defined amount of hydrolases activity for cytokinesis. We propose that modulation of the export of PGHs by STK-dependent signaling pathway is one mechanism of cell adaptation during cell division. As observed in other firmicutes (Hardt et al., 2017; Kaur et al., 2019; Squeglia et al., 2011), PrkC of *C. difficile* could sense extracellular signals generated during PG synthesis (muropeptides, lipid II or other) via its PASTA domains. In WT strain, appropriate hydrolytic activity is controlled by PrkC and STP that can phosphorylate and dephosphorylate CwlA, respectively, limiting or increasing its export as required. In the Δ*prkC* mutant, the non-phosphorylated form of CwlA increases and is more efficiently exported to the CW triggering cell elongation. In the *CD2148* mutant, PrkC could phosphorylate CwlA more efficiently and on multiple sites as observed *in vivo* (Supplementary Fig. 6a), limiting the export of CwlA and resulting in a cell separation defect (**Fig. 8**). In the *CD2148* mutant, the overexpression of CwlA restores a WT phenotype by increasing the amount of non-phosphorylated CwlA able to reach the CW. However, further studies are required to understand the complex regulatory role of CD2148. One hypothesis is that CD2148 might interact at septum of the cells either directly with PrkC or indirectly through proteins of the divisome. These interactions could stimulate PrkC activity or control the choice of its substrates. A second hypothesis is that CD2148 could have a phosphatase activity, thereby desphosphorylating CwlA to increase its export.

**Figure 8.**
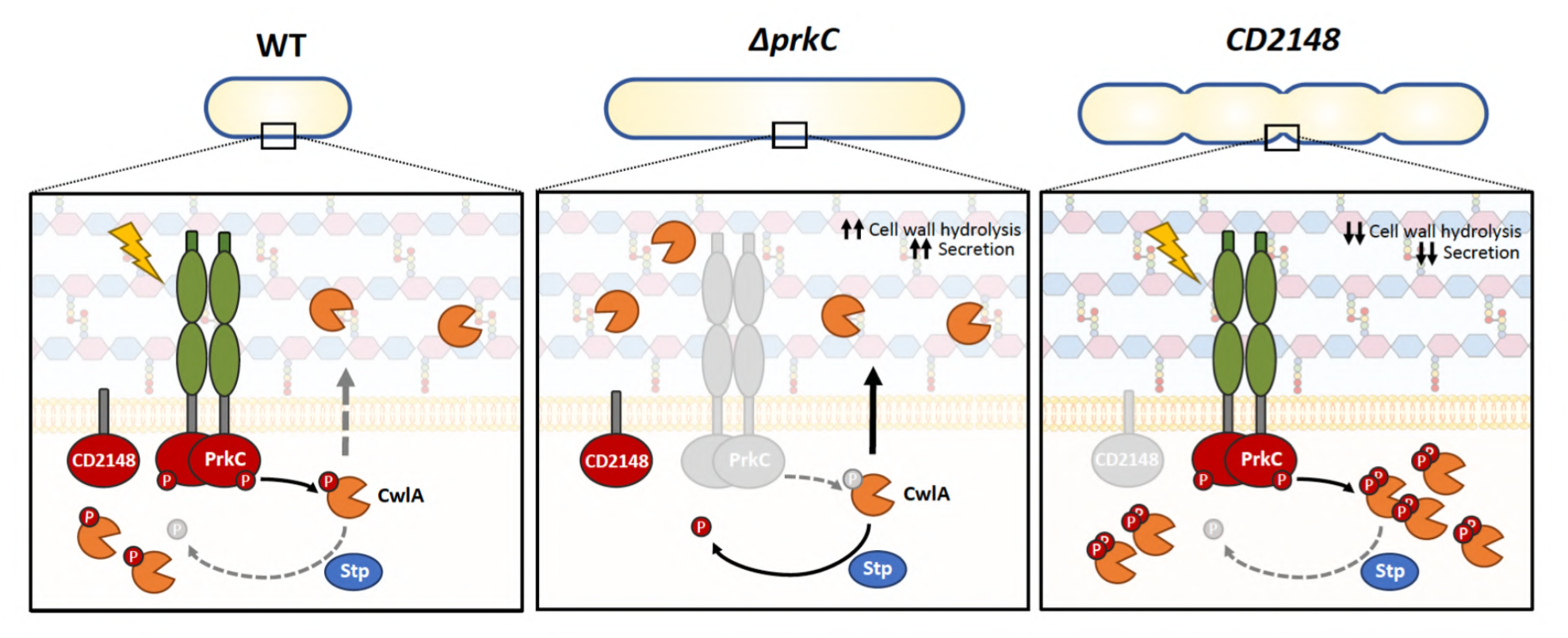
Model for STK-dependent regulation of the endopeptidase CwlA. PrkC could sense extracellular signals generated during PG synthesis (muropeptides, lipid II or others) through its PASTA domains. Active PrkC phosphorylates CwlA (orange scissors) inhibiting its export. Hence, the phosphatase, STP dephosphorylates the PrkC-target CwlA and this protein is exported to the CW, increasing PG hydrolysis when required (*left panel*). In a Δ*prkC* mutant, the non-phosphorylated form of CwlA is more efficiently exported to the CW and high levels of CwlA triggers cell elongation (*middle panel*). In a *CD2148* mutant, PrkC highly phosphorylates CwlA. In consequence, this endopeptidase is less exported resulting in a cell separation defect (*right panel*).

## Methods

### Bacterial strains and growth conditions

Bacterial strains and plasmids used in this study are listed in Supplementary Table 1. *C. difficile* strains were routinely grown at 37°C in an anaerobic environment (90% N_2_, 5% CO_2_, and 5% H_2_) in TY (Bacto tryptone 30 g/L, yeast extract 20 g/L, pH 7.4) or in BHI (Difco). When necessary, *C. difficile* culture media were supplemented with cefoxitin (Cfx; 25 μg/L), cycloserine (Ccs; 250 μg/L), thiamphenicol (Tm; 7.5 μg/L), and erythromycin (Erm; 5 μg/L). *E. coli* strains were cultured at 37°C in LB broth, containing chloramphenicol (25 μg/L), kanamycin (25 μg/ml) or ampicillin (100 μg/L) when necessary. ATc was used to induce expression of the *cwlA* gene from the P_*tet*_ promoter of pDIA6103 (Fagan & Fairweather, 2011). To determine Minimal Inhibitory Concentration (MIC), cultures of *C. difficile* strains (OD_600nm_ of 0.3) were plated on BHI agar plates and the MICs were determined by E-test (bioMérieux) after 24 h incubation at 37°C.

### Construction of *C. difficile* strains and plasmids

All routine plasmid constructions were carried out using standard procedures. To generate the *cwlA::erm* and *CD2148::erm* mutants, the ClosTron system was used as previously described (Heap et al., 2007). Briefly, primers designed to retarget the group II intron on pMTL007 to *cwlA* and *CD2148* were used with the EBS universal primer and intron template DNA to generate a 353-bp DNA fragment for each gene by overlap PCR (Supplementary Table 2). The PCR products were cloned into the HindIII and BsrGI restriction sites of pMTL007 and the sequence of the insertions was verified by sequencing (Supplementary Table 2). Plasmids pMTL007::*cwlA*-1164s and *pMTL::CD2148-302a* retargeted the group II intron for insertion into *cwlA* and *CD2148* in the sense orientation immediately after the 1164th and the 302th nucleotide in the coding sequence, respectively. *E. coli* HB101 (RP4) containing these plasmids were transferred by conjugation into the *C. difficile* 630Δ*erm* strain. Transconjugants selected on BHI plates containing Cfx, Ccs, and Tm were plated on BHI agar containing Erm. Ermresistant *C. difficile* colonies corresponded to plasmid loss and insertion of the group II intron into the chromosome (Heap et al., 2007). The insertion of intron into the target gene was verified by PCR. A Δ*stp* knock out mutant was generated using the *codA*-mediate allele exchange method (ACE) (Cartman et al., 2012; Cuenot et al., 2019; Peltier et al., 2015). 1 kb fragments located up- and downstream of *stp* were amplified from 630Δ*erm* genomic DNA (Supplementary Table 2). Purified fragments were then introduced into the pMTLSC7315 ΔMCS by Gilson Assembly^®^ Master Mix (Biolabs) giving the plasmid pDIA6464 (Supplementary Table 1). *E. coli* HB101 (RP4) containing pDIA6464 was mated with *C. difficile* 630Δ*erm* strain. After conjugation, faster growing single-crossover integrants were isolated by serially restreaking on BHI plates supplemented with Cfx and Tm. Double crossover events were obtained by restreaking single crossover integrants on *C. difficile* minimal medium plates supplemented with fluorocytosine (50 μg.ml^-1^).

To complement the *cwlA::erm* and *CD2148::erm* mutants, the *cwlA* and *CD2148* genes with their own promoter were amplified by PCR. Fragments were introduced into pMTL84121 using the Gibson Assembly^®^ Master Mix. Using *E. coli* HB101 (RP4) as a donor, the resulting plasmids were introduced by conjugation into the *cwlA::erm* or *CD2148::erm* mutants. To construct a plasmid expressing *cwlA* under the control of a P_*tet*_ promoter inducible by ATc, a PCR fragment containing complete gene was amplified and cloned into pDIA6103. For truncated *cwlA* lacking the catalytic NlpC/P60 domain, plasmid pDIA6103-P_*tet*_-*cwlA* was amplified by inverse PCR. The PCR product was then digested by DpnI to remove the plasmid template, phosphorylated by T4 Polynucleotide Kinase and ligated by T4 Ligase to recircularize the plasmid. The same strategy was used to construct the translational fusion coding for a CwlA-HA-tagged protein and to introduce a point mutation into the *cwlA* gene (threonine at position 405 replaced by an alanine [T405A] or a glutamate [T405D]). All generated plasmids and primers are listed in supplementary Tables 1 and 2. pDIA7047 carrying a translational *cwlA*-SNAP fusion was obtained by Gibson Assembly. We amplified the SNAP coding sequence fused to a linker in 5’ orientation (GGATCCGCAGCTGCT) using pFT58 as a template, and pDIA6928 (pDIA6103-*cwlA*) was amplified by inverse PCR (Supplementary Table 2). These plasmids were transferred into *C. difficile* strains by conjugation.

### CwlA homology model

The CwlA 3D homology model was generated using the Phyre2 server (Kelley et al., 2015) (http://www.sbg.bio.ic.ac.uk/phyre2/) based on the alignment with <u>c6biqA</u>, <u>c3npfB</u> and <u>c3h41A</u> (83% of the sequence modelled at >90% accuracy) and displayed in EzMol 2.1 (Reynolds et al., 2018) (http://www.sbg.bio.ic.ac.uk/ezmol/). A sequence alignment of *CwlA* with conserved regions of NlpC/P60 domains of *B. cereus* YkfC, *B. subtilis* YkfC, LytF, LytE and CwlS was generated using Clustal Omega (http://www.ebi.ac.uk/Tools/msa/clustalo/) and ESPript 3.0 (Robert & Gouet, 2014) (http://espript.ibcp.fr/ESPript/ESPript/) to identify the catalytic residues in CwlA belonging to the endopeptidase NlpC/P60 family of proteins.

### Protein synthesis and purification

To purify the kinase domain of PrkC or CD2148, DNA sequence encoding the cytosolic part of each protein, was PCR-amplified from genomic DNA using the primer pairs SAT117/SAT118 and SAT119/SAT120, respectively. The region of *cwlA* coding for the CW-associated hydrolase residues 32 to 432 (the N-terminus amino acids corresponding to the signal peptide were omitted) was amplified using the oligonucleotides SAT285 and SAT286. The PCR products digested with BamHI and KpnI were cloned into the expression vector pQE30, giving plasmids pQE30-*prkC*, pQE30-*CD2148* and pQE30-*cwlA*. These plasmids carried genes encoding proteins fused to a His_6_ tag expressed under the control of an IPTG (isopropylthiogalactoside)-inducible promoter. His_6_-tagged proteins were produced in *E. coli* strain M15pRep4. Cultures were grown at 37°C to an OD_600nm_ of 0.4 and the genes induced 3 h in the presence of 1 mM IPTG. Cells were disrupted by sonication and N-terminus His_6_-tagged proteins were purified on Ni-NTA columns (Qiagen) according to manufacturer’s instruction, desalted with PD-10 columns (GE-Healthcare) and stored at −20°C in a buffer containing 50 mM Tris-Cl pH 7.5, 100 mM NaCl and 10% glycerol. Protein concentrations were estimated using the Bradford assay (Bio-Rad) with BSA as standard.

### *In vitro* phosphorylation assay and Phos-Tag

His_6_-CwlA (10 μM) was incubated alone or in combination with His_6_-PrkC or His_6_-CD2148 (1μM) in kinase buffer (50 mM Tris pH 7.5, 5 mM MgCl_2_). The reaction was initiated by the addition of 5 mM ATP, followed by incubation at 37°C for 1.5 h. Reactions were stopped with the addition of SDS-Laemmli sample buffer and proteins were subjected to Phos-tag™ or SDS-PAGE. Phos-Tag™ is based on the functional molecule Phos-Tag that captures phosphate groups(-PO_3_^2^) (Kinoshita et al., 2006; Kusamoto et al., 2019). Two different methods were performed according to the manufacturer’s instruction: Phos-Tag™ Fluorescence and PhosTag™ Acrylamide. For Phos-Tag™ Fluorescence, SDS-PAGE gels were incubated in a solution containing Phos-Tag (Phos-Tag Phosphoprotein Gel Stain, ABP Biosciences), then washed in Phos-Tag Phosphoprotein Destain Solution (ABP Biosciences). Phosphorylated proteins specifically stained were detected using a fluorescence imaging scanner. For PhosTag™ Acrylamide, which is an electrophoresis technique capable of separating phosphorylated and non-phosphorylated forms based on phosphorylation levels, proteins were run on a 12% SDS-PAGE supplemented with 50 μM of Phos-tag™ Acrylamide (AAL-107, Wako) and 100 μM MnCl_2_. After running, the gel was stained with Coomassie Blue.

### In-gel and FASP-based digestion

His_6_-PrkC was used to phosphorylate His_6_-CwlA *in vitro* as described above. The protein mixture was separated by SDS-PAGE and stained with Coomassie blue. The band corresponding to His_6_-CwlA was excised and subjected to tryptic digestion as previously described (Wiśniewski et al., 2009). Resulting peptides were dried in a Speed-Vac and resuspended in 2% acetonitrile (ACN), 0.1% formic acid (FA) prior to LC-MS/MS analysis.

For FASP-based digestion, bacterial pellets were resuspended in 100 mM ammonium bicarbonate (ABC), 50 mM DTT, 4 % SDS, 1% DNase I, 1X protease and phosphatase inhibitors and disrupted by ultrasonic cavitation. Protein digestion was based on the FASP procedure using 30K Amicon Ultra-4 filtration devices (Millipore). Briefly, 4 mg protein lysate was concentrated into the filtration device at 4500 g for 20 min and diluted with 2 mL of exchange buffer (EB: 100mM ABC, 8 M urea). This step was repeated 3 times before adding 1 mL EB containing 5 mM TCEP, 30 mM chloroacetamide, 0.3 % benzonase, 0.1 % DNase I, 1 mM MgCl_2_ during 1.5 h at RT. After one exchange buffer with EB, the resulting concentrate was washed by 3 steps with 50 mM ABC. After overnight incubation with sequencing grade modified trypsin (Promega) ratio 1:100 (enzyme:protein), peptides were collected by centrifugation of the filter.

### Phospho-enrichment

Resulting peptides were desalted with Sep-Pak plus C18 cartridge (Waters) and eluted with 80 % ACN, 0.1 % heptafluorobutyric acid (HFBA, Sigma-Aldrich) then adjusted at 6 % HFBA. TiO_2_ beads (Sachtopore NP beads 5 μM, 300 Å - Huntsman) were resuspended at 20 mg/mL in 30 % ACN, 0.1 % HFBA during 1 h and activated 15 min with 80% ACN / 6% HFBA. Peptide solution was incubated for 30 min at RT with TiO_2_ ratio 10:1 (bead:peptide). Two washes with the same buffer and one with 50 % ACN, 0.1 % HFBA were performed before an elution with 10 % NH_4_OH. pH was neutralized with 20 % FA and enriched peptides were freeze-dried. Finally, phospho-peptides were desalted by stage-tip using C18 Empore disc and dried in a Speed-Vac. Peptides were resuspended in 2 % ACN / 0.1 % FA prior to LC-MS/MS analysis.

### LC-MS/MS analysis

Tryptic peptides from in gel-digestion were analyzed on a Q Exactive Plus instrument (Thermo Fisher Scientific) and phospho-enrichment peptides on a Q Exactive HF instrument (Thermo Fisher Scientific), both instruments coupled with an EASY nLC 1200 chromatography system (Thermo Fisher Scientific). Sample was loaded on an in-house packed 25 cm (for in geldigestion) and 53 cm (phospho-enrichment) nano-HPLC column with C18 resin (1.9 μm particles, 100 Å pore size, Reprosil-Pur Basic C18-HD resin, Dr. Maisch GmbH, Ammerbuch-Entringen, Germany) after an equilibration step in 100 % solvent A (H_2_O, 0.1 % FA). Peptides were first eluted using a 2 to 5 % gradient of solvent B (ACN, 0.1 % FA) during 5 min, then a 5 to 10 % gradient during 20 min, a 10 to 30 % gradient during 70 min and finally a 30 to 60 % gradient during 20 min, all at 300 nL.min-1 flow rates. The instrument method was set up in the data dependent acquisition mode. After a survey scan in the Orbitrap (resolution 70,000 and 60,000), the 10 most intense precursor ions were selected for HCD fragmentation with a normalized collision energy set up to 27 and 28. Charge state screening was enabled, and precursors with unknown charge state or a charge state of 1, 7, 8 and >8 were excluded. Dynamic exclusion was enabled for 20s and 30s.

### Data processing for protein identification and quantification

Raw files were searched Maxquant software (Tyanova et al., 2016) version 1.5.3.8 with Andromeda (Cox et al., 2011) as a search engine against an internal *C. difficile* database containing 3,957 proteins (Cuenot et al., 2019), usual known mass spectrometry contaminants and reversed sequences of all entries. Andromeda searches were performed choosing trypsin as specific enzyme with a maximum number of three missed cleavages. Possible modifications included carbamidomethylation (Cys, fixed), oxidation (Met, variable), Nter acetylation (variable) and phospho (Ser, Thr, Tyr, variable). The mass tolerance in MS was set to 20 ppm for the first search then 4.5 ppm for the main search and 20 ppm for the MS/MS. Maximum peptide charge was set to seven and seven amino acids were required as minimum peptide length.

The “match between runs” feature was applied for samples having the same experimental condition with a maximal retention time window of 0.7 minute. One unique peptide to the protein group was required for the protein identification for the phospho-enrichment analysis. A false discovery rate (FDR) cutoff of 1 % was applied at the peptide and protein levels. For in-gel analysis, we used the Phospho (STY) table to extract the intensity of each phosphopeptide. A normalization step based on the iBAQ of the protein in each sample was performed before relative quantification. The mass spectrometry proteomics data have been deposited to the ProteomeXchange Consortium (http://proteomecentral.proteomexchange.org) via the PRIDE partner (Vizcaino et al., 2014) repository with the dataset identifier PXD021541.

### Detection of PG-hydrolyzing activity and identification of hydrolytic specificity of CwlA

PG samples were prepared from 600 mL of different strains of *C. difficile* grown in TY (OD_600nm_ of 1). PG-PSII was first purified as previously described (Cuenot et al., 2019) and used for the detection of the lytic activity of the purified enzymes by zymogram analysis. *C. difficile* purified PG was resuspended in dH_2_O, and the suspension was added to an SDS-polyacrylamide gel at a final concentration of 1 mg/mL. After electrophoresis, the gel was shaken at 37°C for 16 h in 50 ml of 25 mM Tris-HCl (pH 8.0) solution containing 1 % (vol/vol) Triton X-100 to allow protein renaturation. Clear bands resulting from lytic activity were visualized after staining with 1% (wt/vol) methylene blue (Sigma)-0.01% (wt/vol) KOH and subsequent destaining with distilled water (Fukushima & Sekiguchi, 2016; Wydau-Dematteis et al., 2018).

The PG was further separated from PSII by hydrofluoric acid treatment (Cuenot et al., 2019). To identify the hydrolytic specificity of CwlA, purified PG (0.75 mg) was incubated with 100 μg of purified His_6_-CwlA in 50 mM Tris-HCl, pH 8.0 overnight at 37°C. A control sample of PG was incubated without enzyme in the same conditions. After incubation, soluble and insoluble fractions were separated by centrifugation at 20,000 g for 15 min. The insoluble fraction was further digested with mutanolysin from *Streptomyces globisporus* (Sigma) at 2500 u.mL^-1^ for 19 h at 37°C in 25 mM sodium phosphate under shaking. The soluble muropeptides were reduced with sodium borohydride and analyzed by RP-HPLC and mass spectrometry as described previously (Peltier et al., 2011).

### Phase contrast, SNAP labelling and fluorescence microscopy

For phase contrast microscopy, *C. difficile* strains were cultured for 5 h in TY (with antibiotics and inducers when needed) at 37°C. Cells were visualized using Zeiss Axioskop microscope and analysed using the software ImageJ and the plugin MicrobeJ (Ducret et al., 2016) for quantitative analysis. For membrane staining, 500 μL of exponentially growing cells in TY were centrifuged and resuspended in 100 μL of PBS supplemented with the fluorescent membrane dye FM 4-64 (Molecular Probes, Invitrogen) at 1 μg/mL. Samples were incubated for 2 min in the dark and mounted on 1.2% agarose-coated glass slides. For SNAP labelling, strains were grown 3 h in TY and the expression of the *cwlA*-SNAP^*Cd*^ or SNAP^*Cd*^-*CD2148* fusions was induced with 50 ng/mL of ATc for 2 h. The TMR-Star substrate (New England Biolabs) was added at 250 nM, and the mixture was incubated for 30 min in the dark under anaerobiosis. Cells were then collected by centrifugation, washed and resuspended in PBS. Cell suspension (3 μL) was mounted on 1.2 % agarose pad. The images were taken with exposure times of 600 ms for autofluorescence and 800 ms for SNAP using a Nikon Eclipse TI-E microscope 100x Objective and captured with a CoolSNAP HQ2 Camera. The images were analysed using ImageJ.

### Transmission electron microscopy (TEM)

*C. difficile* strains were grown in TY at 37°C for 5 h. After centrifugation at 5000 rpm for 10 min at 4°C, cells pellets were resuspended in 2% glutaraldehyde in 0.1 M cacodylate buffer and incubated 1 h at room temperature. Then, cells pellets were washed with 0.2 M sucrose in 0.1 M cacodylate buffer. Staining and examinations were done by the GABI-MIMA2 TEM Platform at INRA, Jouy-en-Josas, France. Grids were examined with a Hitachi HT7700 electron microscope operated at 80 kV, and images were acquired with a charge coupled device camera (AMT).

### Cell lysis, fractionation, and protein analysis

We confirmed the production of a CwlA-HA-tagged protein by Western blot using an antibody raised against the HA tag. Cellular fractions were extracted as previously described (Peltier et al., 2015). Briefly, *C. difficile* cultures (10 mL) were harvested by centrifugation at 5,000 ×g for 10 min at 4°C, culture supernatants (Sn) were filtered through a 0.22-μm-pore-size filter and precipitated on ice with 10% TCA for 30 min (Sn) prior to SDS-PAGE. Pellets resuspended in phosphate-sucrose buffer (0.05 M HNa_2_PO_4_, pH 7.0, 0.5 M sucrose) to an OD_600nm_ of 40 were incubated at 37 °C for 1 h in the presence of purified CD27L endolysin (30 μg/mL). The protoplasts were then recovered by centrifugation at 6000 ×g for 20 min at 4 °C. Supernatants containing the cell wall (CW) fraction were removed, and the protoplast pellet was resuspended in phosphate buffer (0.05 M HNa_2_PO_4_, pH 7.0) containing 40 μg/mL DNase I at an OD_600nm_ of 40 and incubated at 37 °C for 45 min. Lysates were harvested at 16,000 ×g for 15 min at 4 °C to separate supernatants containing the cytoplasmic (Cyto) fraction and the membrane pellet. For analysis by SDS-PAGE, an equal volume of 2 X SDS sample buffer was added to protein samples. SDS-PAGE and Western immunoblotting were carried out using standard methods. Proteins were electrophoresed and transferred to polyvinylidene difluoride membranes (PVDF). After blocking with non-fat milk in TBST buffer, primary antibodies were added. The washed membranes were incubated with appropriate secondary antibodies coupled to horseradish peroxidase that were detected by an enhanced chemiluminescence system. Antibodies against the HA and SNAP epitope were purchased from Osenses and New England BioLabs, respectively.

## Acknowledgements

This work was funded by the Institut Pasteur, the Université de Paris and by grants from the ITN Marie Curie, Clospore (H2020-MSCA-ITN-2014 642068) and the ANR DifKin (ANR-17-CE15-0018-01). EC and TGG are the recipient of a ITN Marie Curie and an ANR fellowships, respectively. The anti-Cwp66 antibody was a gift from N. Fairweather. This work has benefited from the facilities and expertise of Christine Longin and MIMA2 MET, INRA-UMR GABI.

## Author contributions

TGG, TC and IMV conceived and designed the study, TGG, SP, EC, TD, JP, PC performed the experiments, TGG, MPCC, MM, QGG, BD, TC, IMV analyzed the data, TGG, TC and IMV wrote the manuscript. All authors revised and commented on the manuscript.

---

**Supplementary Figure 1.**
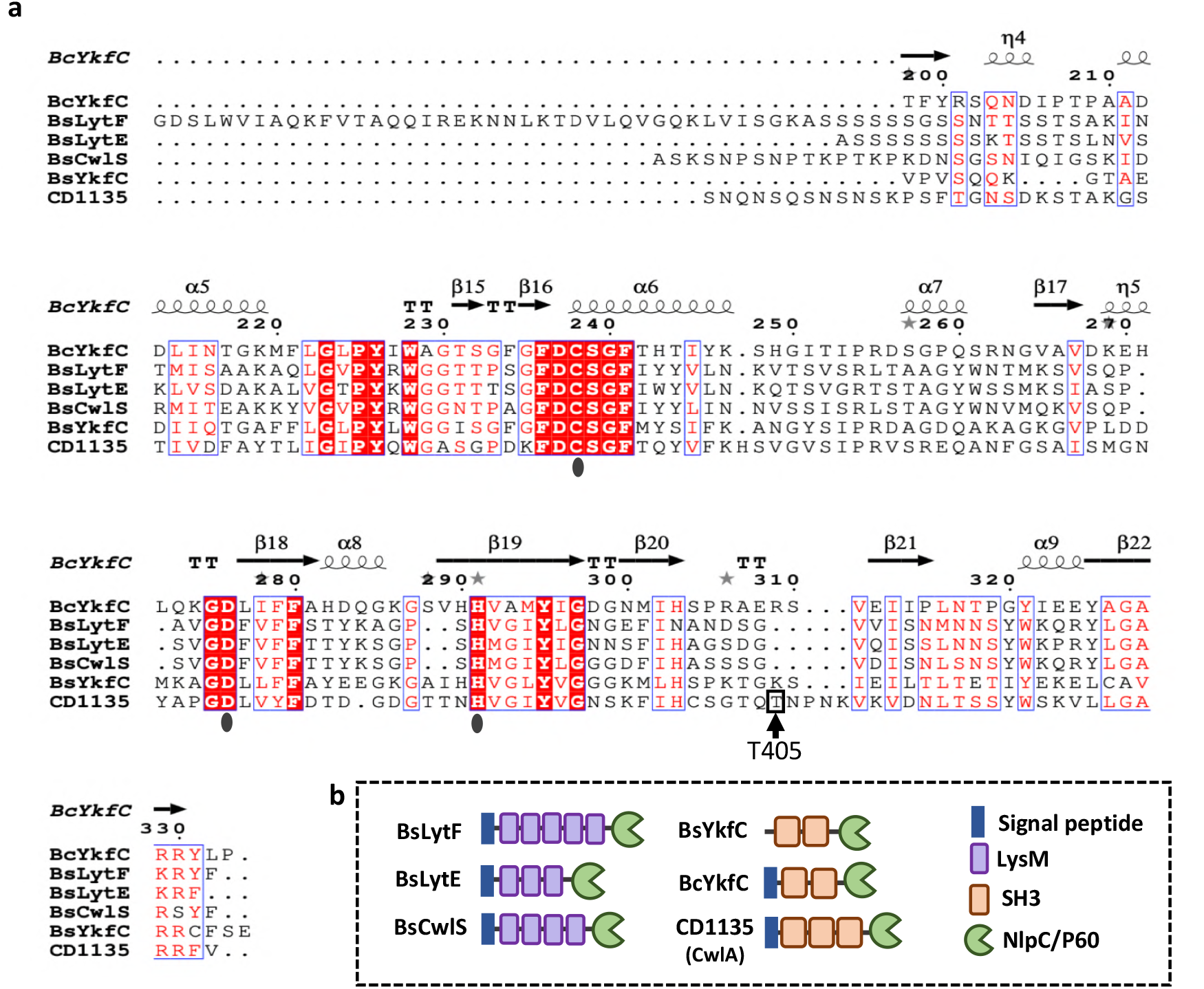
CD1135 (CwlA) belongs to the NlpC/P60 family of endopeptidases. **a,** Alignment of CD1135 with conserved regions of NlpC/P60 domains of *B. cereus* YkfC (BcYkfC), *B. subtilis* YkfC (BsYkfC), LytF (BsLytF), LytE (BsLytE) and CwlS (BsCwlS) contain the essential residues for catalysis (Cys, His, Asp; point black). Conserved residues are in red boxes and similar residues in red characters. **b,** Schematic representation of *B. subtilis* LytF, LytE, CwlS, YkfC, *B. cereus* YkfC and CD1135.

**Supplementary Figure 2.**
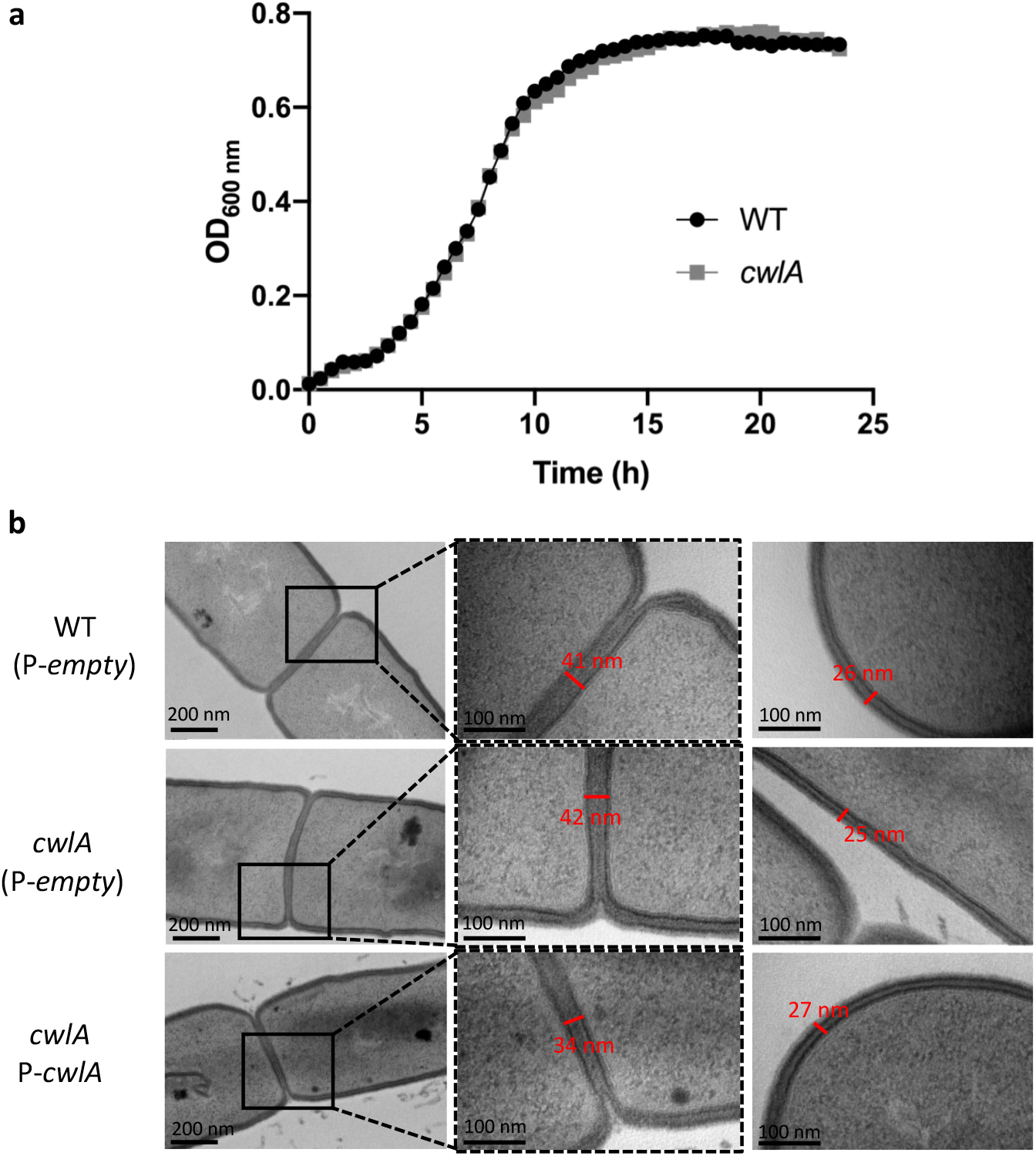
Growth and TEM of the *cwlA* mutant. **a,** Growth curves of the *cwlA* mutant compared to WT strain in TY. **b,** Transmission electron microscopy showing septa thickness and cell wall width of 630Δ*erm* + P-*empty* (WT), *cwlA* + P-*empty* and *cwlA* + P-*cwlA*. Middle and right columns: higher magnifications (100 nm) of the division septa (highlighted by a black square) and cell wall, respectively.

**Supplementary Figure 3.**
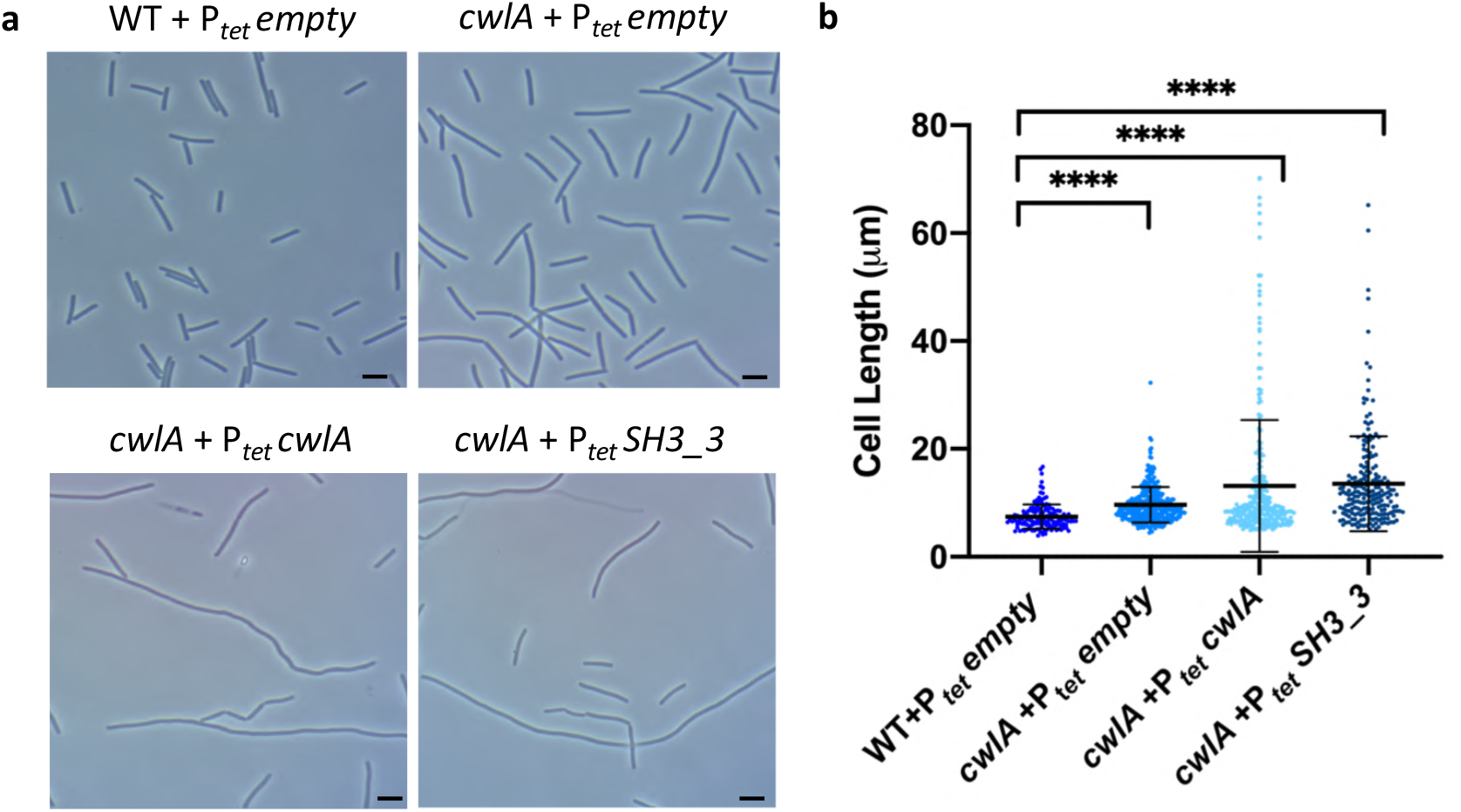
Morphology of cells overexpressing *cwlA*. **a**, Phase contrast images of *C. difficile* cells of WT + P_*tet*_ *empty, cwlA* + P_*tet*_ *empty, cwlA* + P_*tet*_ *cwlA* and *cwlA* + P_*tet*_ *SH3_3*. Scare bar, 5 μm. **b**, Scatter plots showing the distribution of cell length. Two-sided Mann–Whitney *U* tests (\*\*\*\**P* < 0.0001), cells counted 148 (WT + P_*tet*_ *empty*), 339 (*cwlA* + P_*tet*_ *empty*), 290 (*cwlA* + P_*tet*_ *cwlA*) and 208 (*cwlA* + P_*tet*_ *SH3_3*) in a single representative experiment. Experiments were performed in triplicate.

**Supplementary Figure 4.**
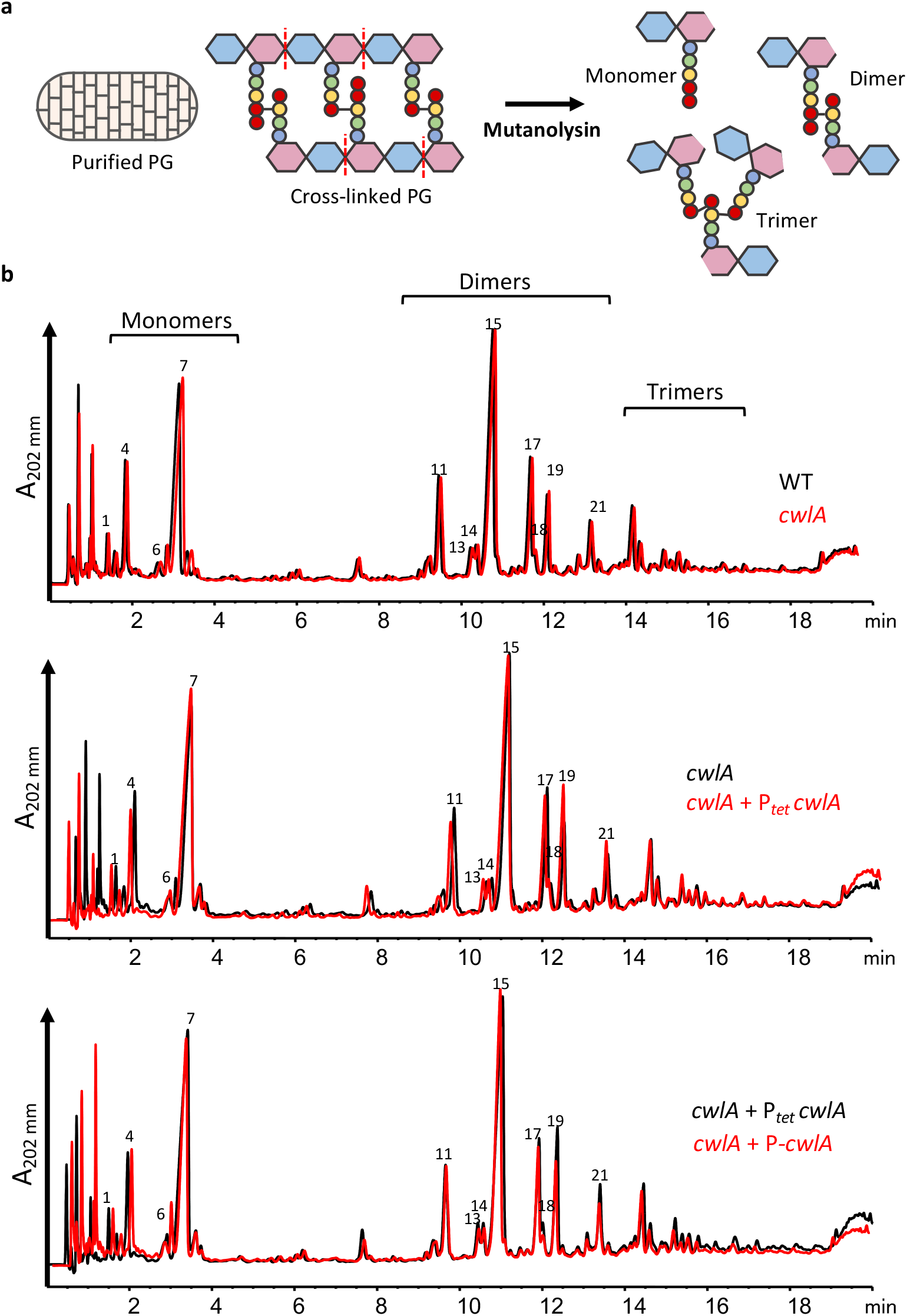
Profiles of muropeptides of *cwlA* strains. **a,** Schematic representation of PG isolation and digestion for RP-HPLC analysis, mutanolysin treatment of cross-linked PG generate muropeptides species. **b,** RP-HPLC separation profile of muropeptides from WT (P_*tet*_ *empty*), *cwlA* (P_*tet*_ *empty*), *cwlA* + P_*tet*_ *cwlA* and *cwlA* + P-*cwlA* (*cwlA + pMTL84121-cwlA*). The profiles were superimposed and the peaks were numbered according to Cuenot et al., 2019. Peak numbers 1, 4, 6, 7 corresponds to monomers and peak numbers 11, 13, 14, 15, 17, 18, 19 and 21 corresponds to dimers. The abundance of each molecule is correlated to the area present under the corresponding peak

**Supplementary Figure 5.**
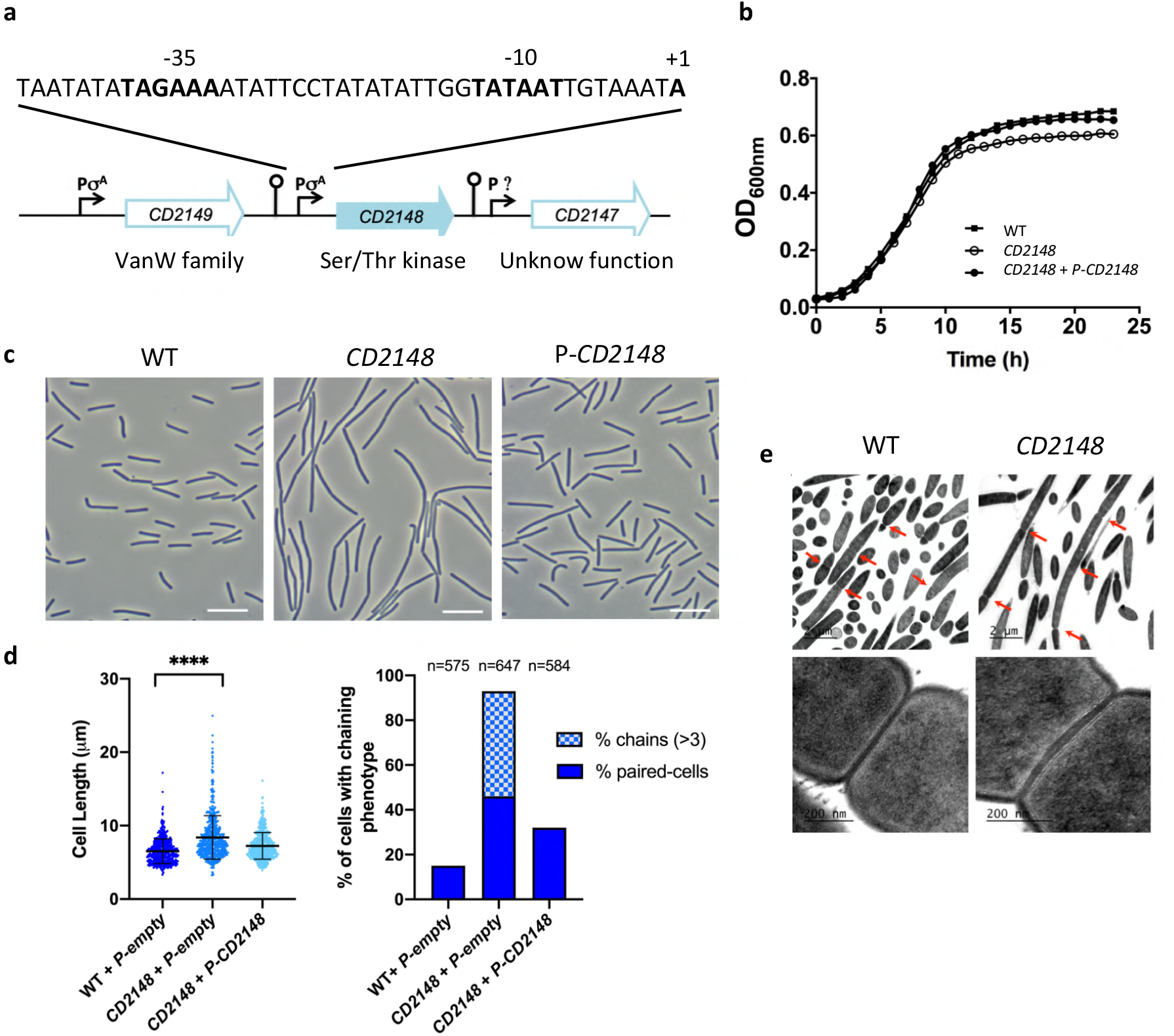
Functional characterization of CD2148. **a**, Genetic organization of the *CD2148* locus. TSS mapping experiment indicated the presence of a σ^A^-dependent promoter upstream of *CD2148* and *CD2149* (Soutourina et al., 2020). The −35 and −10 boxes as well as the TSS are indicated in bold. **b**, Growth curves of *CD2148* mutant compared to WT strain and the complemented strain *CD2148* + P-*CD2148* in TY. **c**, Phase contrast images of WT (630 Δ*erm* + pMTL84121), *CD2148* (*CD2148::erm* + pMTL84121) and *CD2148* + P-*CD2148* (*CD2148::erm* + pMTL84121-*CD2148*) cells grown in TY during exponential phase. **d**, Scatter plots showing cell length (left) and percentage of cells harboring a chaining phenotype (right). *P* values were determined by two-sided Mann–Whitney *U* tests (\*\*\*\**P* < 0.0001); counted 575 (WT + P-*empty*), 647 (*CD2148* + P-*empty*) and 584 (*CD2148* + P-*CD2148*). **e**, Transmission electron microscopy showing division septa (red arrows, *upper panel*), scale bars 2 μm. Higher magnifications (200 nm) of septa thickness (*lower panel*).

**Supplementary Figure 6.**
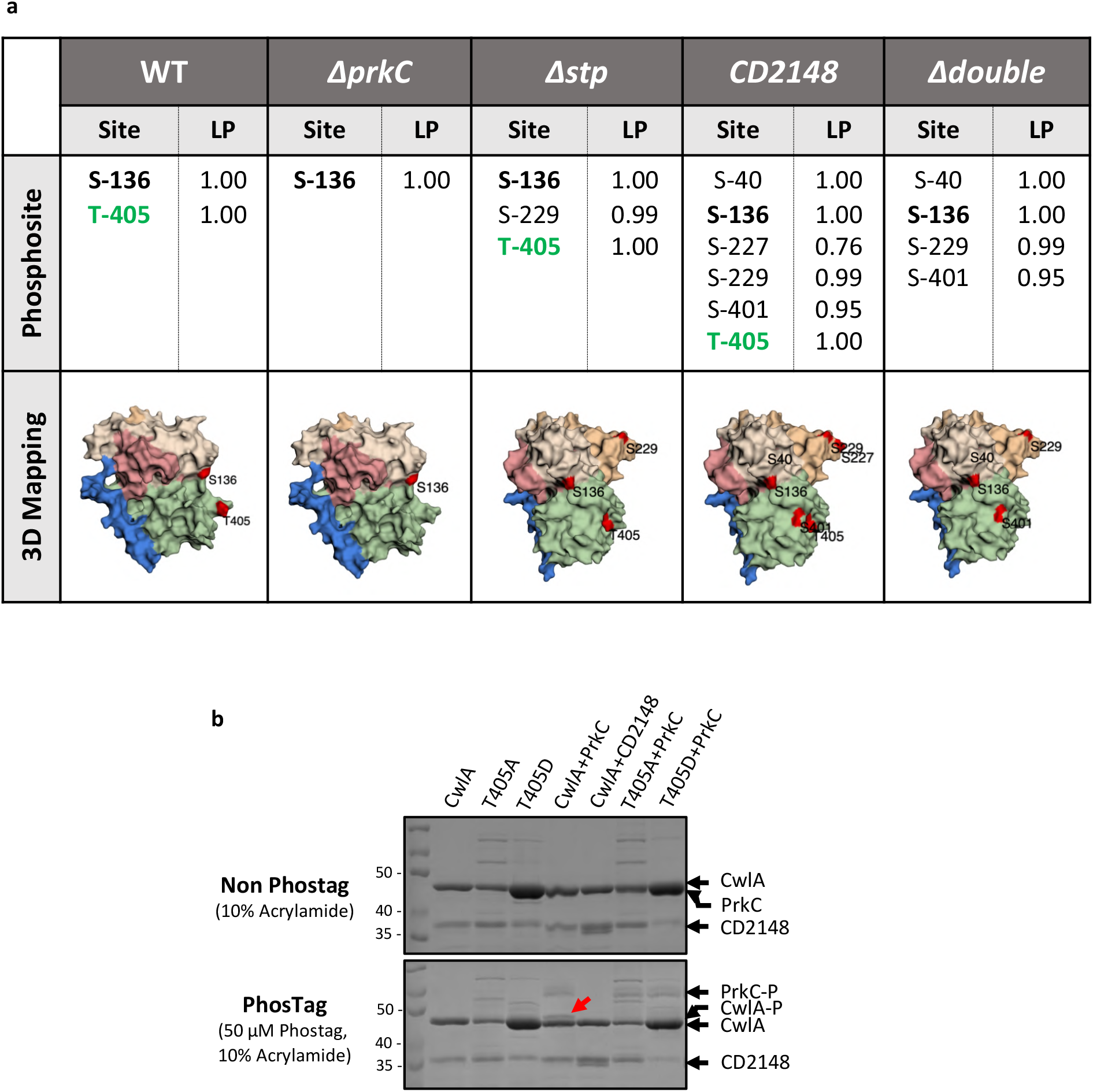
*In vivo* and *in vitro* phosphorylation of CwlA. **a,** Surface representation of CwlA showing phosphorylated residues identified *in vivo* in the STKs/STP mutants with a good localization probability (LP>0.75). Phosphosites S-136 and T-405 are highlighted in black and green, respectively. CwlA is colored by domains (blue, signal peptide; pink, SH3_3.1; light pink, SH3_3.2; orange, SH3_3.3 and green, NlpC/P60) and phosphosites are labeled in red. **b,** *In vitro* phosphorylation assay of CwlA, CwlA-T405A and CwlA-T405D by PrkC or CD2148 and visualized by Phos-tag™ Acrylamide. Red arrow indicate phosphorylated CwlA.

**Supplementary Figure 7.**
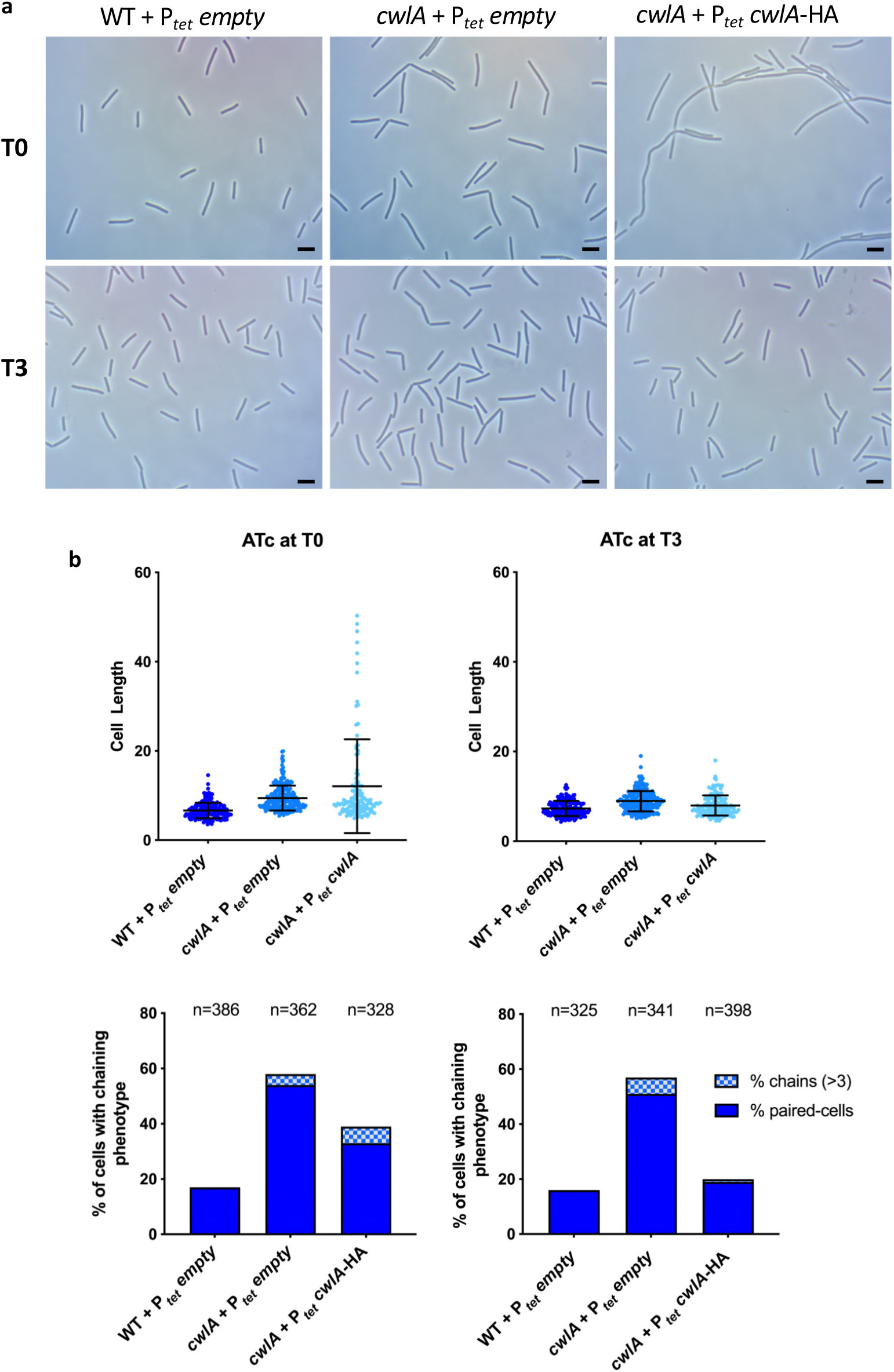
Complementation of *cwlA* mutant with pDIA6103-P_*tet*_ *cwlA* plasmid. **a,** Phase contrast images of WT, *cwlA* and *cwlA*+ P_*tet*_ *cwlA* cells after 5 h of growth in TY. Induction performed with 50 ng/ml ATc from the beginning of inoculation (T0) or after 3 h of growth (T3). Scare bar, 5 μm. **b**, Scatter plots showing cell length with the median and SD of each distribution indicated by a black line (*upper*) and percentage of cells harboring a chaining phenotype (*lower*), n indicates the number of cells counted per strain in a single representative experiment. The images and n values are representatives of experiments performed in triplicate.

**Supplementary Figure 8.**
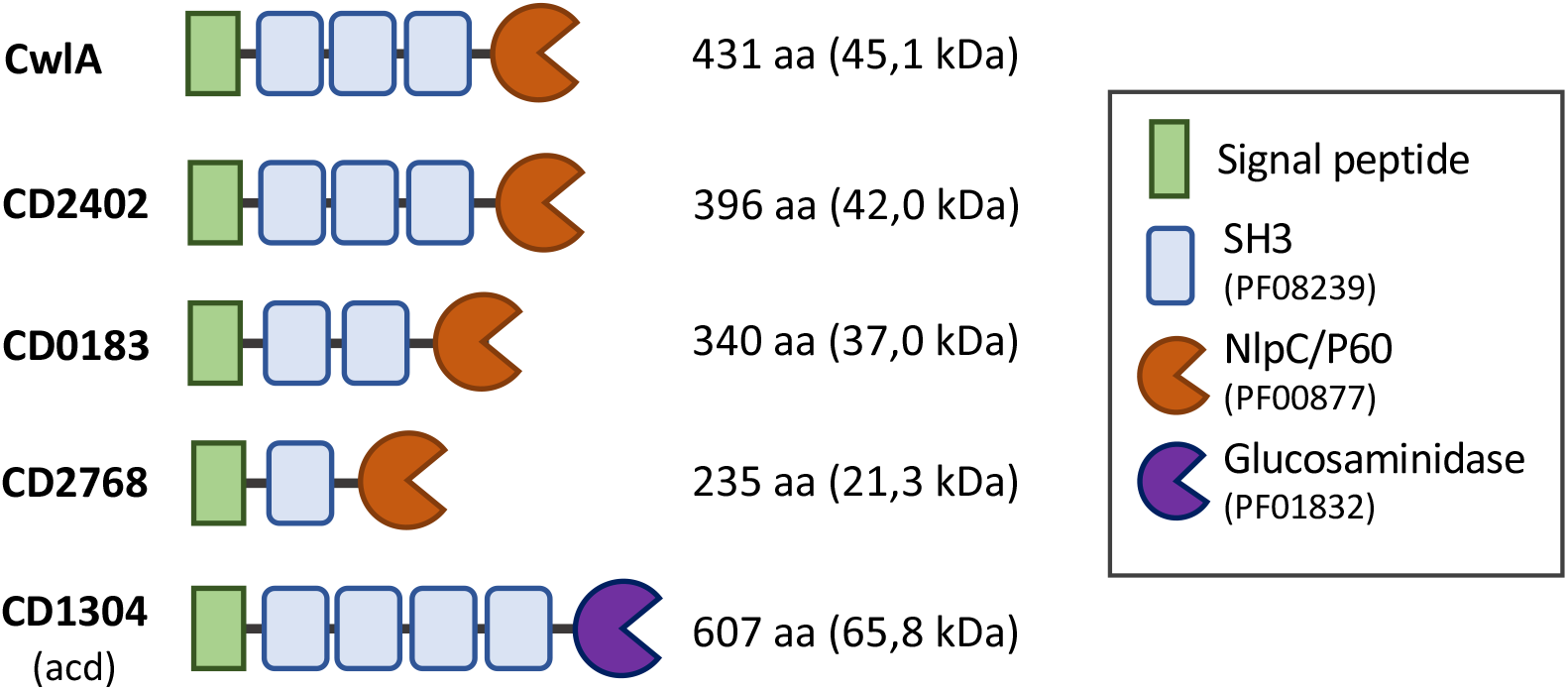
Predicted PG-degrading enzymes associated to SH3 domains identified in *C. difficile* 630. Schematic representation of PG-degrading enzymes containing different numbers of SH3 domains (PF08239).

## Supplementary Tables

**Supplementary Table 1.** Strains and plasmids used in this study

| *E. coli* strains |  |  |
| --- | --- | --- |
| NEB |  |  |
| HB101 (RP4) | *supE44 aa14 galK2 lacY1 Δ (gpt-proA) 62 rpsL20 (Str<sup>R</sup>)xyl-5 mtl-1 recA13 Δ (mcrC-mrr) hsdS<sub>B</sub>(r<sub>B</sub><sup>-</sup>m<sub>B</sub><sup>-</sup>) RP4 (Tra<sup>+</sup> IncP Ap<sup>R</sup> Km<sup>R</sup> Tc<sup>R</sup>)* | Laboratory stock |
| BL21 | *F<sup>-</sup> ompT gal dcm lon hsdS<sub>B</sub>(r<sub>B</sub><sup>-</sup>m<sub>B</sub><sup>-</sup>) λ(DE3 [lacI lacUV5-T7 gene 1 ind1 sam7 nin5])* | Novagen |
| M15pRep4 |  |  |
| *C. difficile* strains |  |  |
| 630Δ*erm* | wild-type | Laboratory stock |
| CDIP631 | 630Δ*erm* Δ*prkC* | Cuenot *et al.*, 2019 |
| CDIP823 | 630Δ*erm* Δ*stp* | This work |
| CDIP736 | 630Δ*erm* CD2148::*erm* | This work |
| CDIP738 | 630Δ*erm* Δ*prkC* CD2148::*erm* | This work |
| CDIP1150 | 630Δ*erm* *cwlA*::*erm* | This work |
| CDIP892 | 630Δ*erm* (pMTL84121) | Laboratory stock |
| CDIP1735 | 630Δ*erm* *cwlA*::*erm* (pMTL84121) | This work |
| CDIP893 | 630Δ*erm* CD2148::*erm* (pMTL84121) | This work |
| CDIP1665 | 630Δ*erm* *cwlA*::*erm* pMTL84121-*cwlA* | This work |
| CDIP894 | 630Δ*erm* CD2148::*erm* pMTL84121-CD2148 | This work |
| CDIP219 | 630Δ*erm* (pDIA6103) | Laboratory stock |
| CDIP1548 | 630Δ*erm* *cwlA*::*erm* (pDIA6103) | This work |
| CDIP1549 | 630Δ*erm* *cwlA*::*erm* pDIA6103 P<sub>tet</sub>-*cwlA* | This work |
| CDIP1643 | 630Δ*erm* *cwlA*::*erm* pDIA6103 P<sub>tet</sub>-*cwlA*-HA | This work |
| CDIP1644 | 630Δ*erm* *cwlA*::*erm* pDIA6103 P<sub>tet</sub>-*cwlA* T405A-HA | This work |
| CDIP1645 | 630Δ*erm* *cwlA*::*erm* pDIA6103 P<sub>tet</sub>-*cwlA* T405D-HA | This work |
| CDIP1474 | 630Δ*erm* pDIA6103 P<sub>tet</sub>-*cwlA*-HA | This work |
| CDIP1475 | 630Δ*erm* Δ*prkC* pDIA6103 P<sub>tet</sub>-*cwlA*-HA | This work |
| CDIP1476 | 630Δ*erm* CD2148::*erm* pDIA6103 P<sub>tet</sub>-*cwlA*-HA | This work |
| CDIP1477 | 630Δ*erm* Δ*stp* pDIA6103 P<sub>tet</sub>-*cwlA*-HA | This work |
| CDIP1646 | 630Δ*erm* Δ*prkC* CD2148::*erm* pDIA6103 P<sub>tet</sub>-*cwlA*-HA | This work |
| CDIP1660 | 630Δ*erm* *cwlA*::*erm* P<sub>tet</sub>-*cwlA*-SNAP<sup>Cd</sup> | This work |
| CDIP1659 | 630Δ*erm* CD2148::*erm* P<sub>tet</sub>-SNAP<sup>Cd</sup>-CD2148 | This work |
| CDIP1357 | 630Δ*erm* P<sub>tet</sub>-SNAP<sup>Cd</sup>-*prkC* | Cuenot *et al.*, 2019 |
| Plasmids |  |  |
| pMTL007 | group II intron, ErmBtdRAM2 and ltrA ORF from pMTL20lacZTTErmBtdRAM2 Cm<sup>R</sup> | Heap *et al.*, 2007 |
| pMTL007::*cwlA* | pMTL007::*cwlA*-1164s | This work |
| pDIA6485 | pMTL007::CD2148-302a | This work |
| pDIA6464 | pMTLSC7315 Δ*stp* | This work |
| pMTL84121 | *Clostridia* modular plasmid; *catP* (Cm<sup>R</sup>/Tm<sup>R</sup>) | Heap *et al.*, 2009 |
| pDIA6103 | pRPF185 Δ*gusA* | Soutourina *et al.*, 2013 |
| pDIA6912 | pMTL84121-*cwlA* | This work |
| pDIA6712 | pMTL84121-*CD2148* | This work |
| pDIA6928 | pDIA6103-P<sub>tet</sub>-*cwlA* | This work |
| pDIA6935 | pDIA6103-P<sub>tet</sub>-*cwlA*-HA | This work |
| pDIA6991 | pDIA6103-P<sub>tet</sub>-*cwlA* -T405A (phospho-ablative) | This work |
| pDIA6993 | pDIA6103-P<sub>tet</sub>-*cwlA*-T405D (phospho-mimetic) | This work |
| pDIA7026 | pDIA6103-P<sub>tet</sub>-*cwlA*-T405A-HA (phospho-ablative) | This work |
| pDIA7027 | pDIA6103-P<sub>tet</sub>-*cwlA*-T405D-HA (phospho-mimetic) | This work |
| pDIA7047 | pDIA6103-P<sub>tet</sub>-*cwlA*-SNAP<sup>Cd</sup> | This work |
| pDIA7046 | pDIA6103-P<sub>tet</sub>-SNAP<sup>Cd</sup>-*CD2148* | This work |
| pDIA6855 | pDIA6103-P<sub>tet</sub>-SNAP<sup>Cd</sup>-*prkC* | Cuenot *et al.*, 2019 |
| pDIA6406 | pQE30-*prkC* | This work |
| pDIA6407 | pQE30-*CD2148* | This work |
| pDIA7091 | pQE30-*cwlA* | This work |
| pDIA7018 | pQE30-*cwlA* T405A | This work |
| pDIA7019 | pQE30-*cwlA* T405D | This work |

**Supplementary Table 2.**
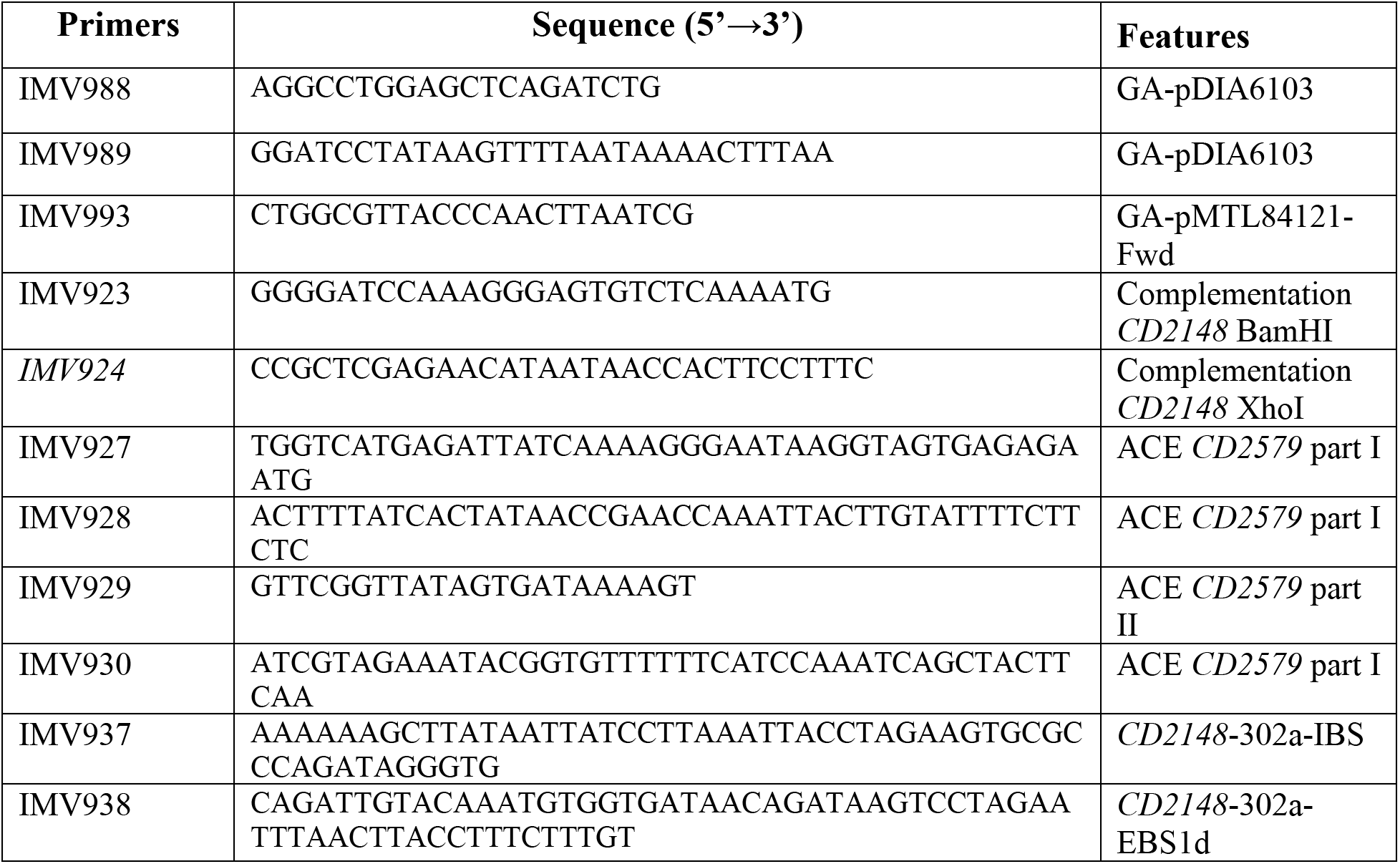
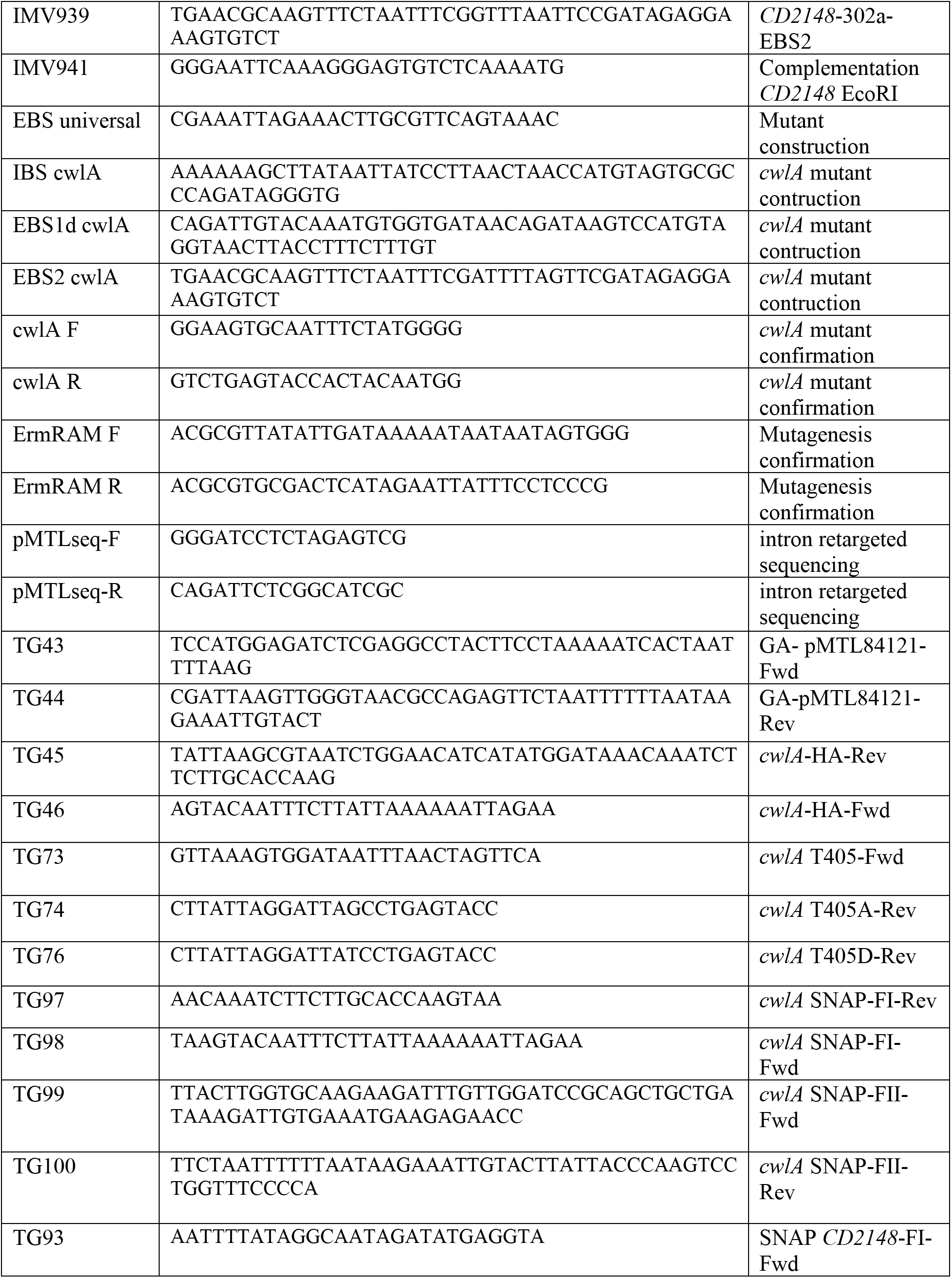
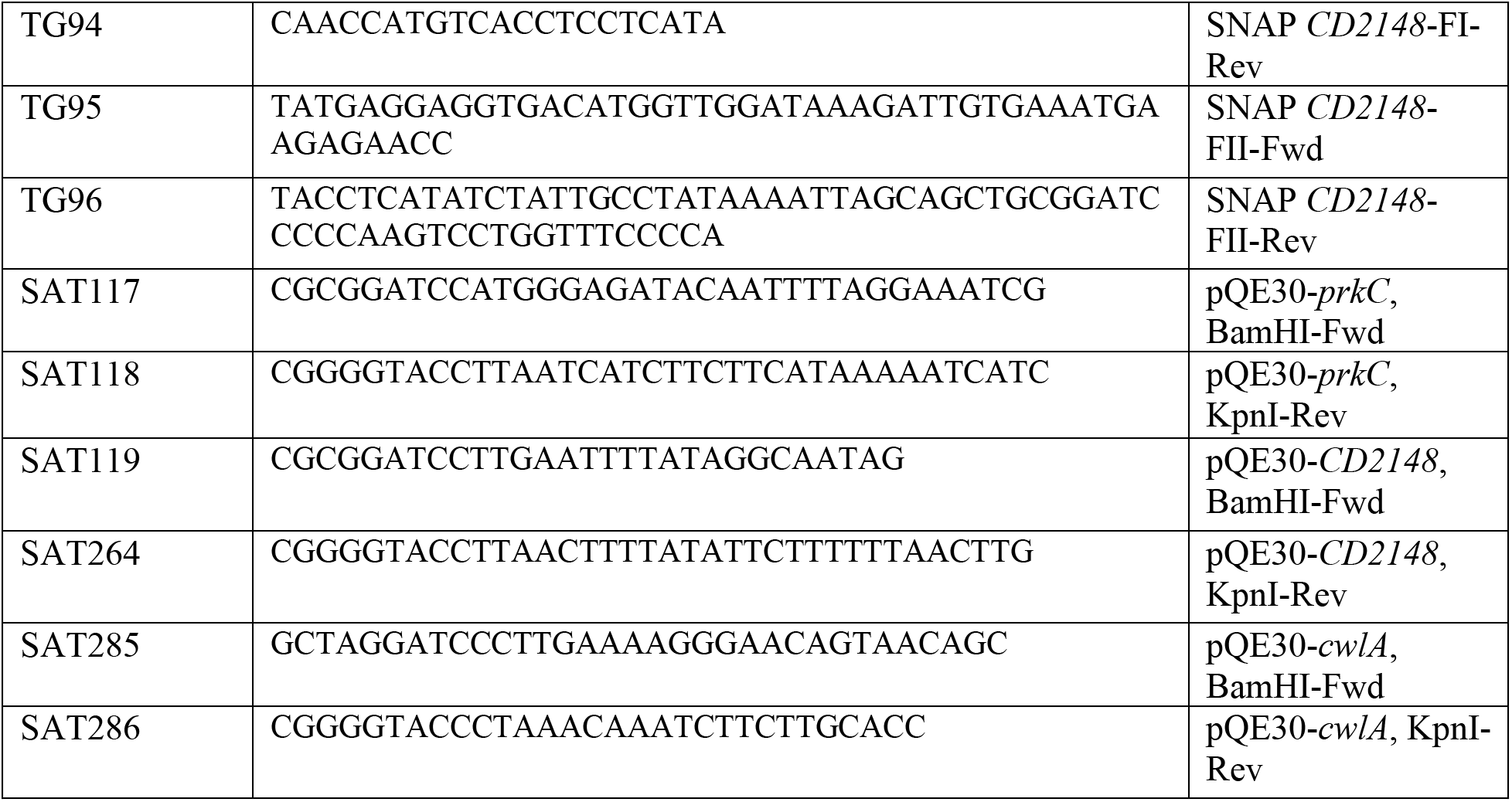
Oligonucleotides used in this study

